# Integrating multiple experimental data to determine conformational ensembles of an intrinsically disordered protein

**DOI:** 10.1101/2020.02.05.935890

**Authors:** Gregory-Neal W. Gomes, Mickaël Krzeminski, Ashley Namini, Erik. W. Martin, Tanja Mittag, Teresa Head-Gordon, Julie D. Forman-Kay, Claudiu C. Gradinaru

## Abstract

Intrinsically disordered proteins (IDPs) have fluctuating heterogeneous conformations, which makes structural characterization challenging, but of great interest, since their conformational ensembles are the link between their sequences and functions. An accurate description of IDP conformational ensembles depends crucially on the amount and quality of the experimental data, how it is integrated, and if it supports a consistent structural picture. We have used an integrative modelling approach to understand how conformational restraints imposed by the most common structural techniques for IDPs: Nuclear Magnetic Resonance (NMR) spectroscopy, Small-angle X-ray Scattering (SAXS), and single-molecule Förster Resonance Energy Transfer (smFRET) reach concordance on structural ensembles for Sic1 and phosphorylated Sic1 (pSic1). To resolve apparent discrepancies between smFRET and SAXS, we integrated SAXS data with non-smFRET (NMR) data and reserved the new smFRET data for Sic1 and pSic1 as an independent validation. The consistency of the SAXS/NMR restrained ensembles with smFRET, which was not guaranteed a priori, indicates that the perturbative effects of NMR or smFRET labels on the Sic1 and pSic1 ensembles are minimal. Furthermore, the mutual agreement with such a diverse set of experimental data suggest that details of the generated ensembles can now be examined with a high degree of confidence to reveal distinguishing features of Sic1 vs. pSic1. From the experimentally well supported ensembles, we find they are consistent with independent biophysical models of Sic1’s ultrasensitive binding to its partner Cdc4. Our results underscore the importance of integrative modelling in calculating and drawing biological conclusions from IDP conformational ensembles.

## 1 Introduction

Under physiological conditions, the amino acid sequences of intrinsically disordered proteins (IDPs) encode for a large and heterogeneous ensemble of conformations, allowing them to perform critical biological functions. ^1,2^ The properties of IDP conformational ensembles are intimately related to their function in health and disease. ^3^ This has prompted intense efforts to develop formal and heuristic descriptions of how sequence properties relate to conformational ensembles, ^4–8^ and how the properties of conformational ensembles, once determined, can be mined to generate hypotheses about biological function. ^9–11^ Conformational ensembles are therefore central to understanding both sequence-to-ensemble and ensemble-to-function relationships in IDPs, which makes their accurate and comprehensive characterization of high importance.

To provide insights into the structural properties of IDPs, Nuclear Magnetic Resonance (NMR), ^12^ Small-Angle X-Ray Scattering (SAXS), ^13^ and single-molecule Förster Resonance Energy Transfer (smFRET) ^14,15^ have emerged as particularly powerful techniques. Computational approaches to integrate the information from these measurements typically represent conformational ensembles as a collection of structures, each described by its atomic coordinates, and use the experimental data for constructing (e.g., restraining or re-weighting), or validating the ensemble calculation. ^16–18^

Despite their demonstrated complementarity, ^19–21^ conformational ensembles which use data from all three techniques in their construction or validation are rarely reported. Aznauryan et al., reported ensembles of ubiquitin denatured in 8 M urea which are consistent with SAXS and a large number of restraints from NMR and smFRET experiments. ^20^ However, concerns about the mutual consistency of smFRET and SAXS data posit that in the absence of denaturant, the FRET fluorophores could interact with each other and/or the IDP itself. ^22^ Piana et al., reported ensembles of *α*-synuclein in physiological conditions, which are directly compared to SAXS and NMR data, but are compared to distances inferred from smFRET data using an assumed homopolymer model. ^23^ However, it is difficult to determine which, if any, homopolymer model is appropriate for a particular heteropolymeric IDP. ^24,25^ Thus, using data from all three techniques to construct or validate conformational ensembles of an IDP (*i*) in *physiological* conditions and (*ii*) without assuming a homopolymer model, would provide valuable insights into each technique’s sensitivity to different aspects of IDP structure.

We therefore sought to determine conformational ensembles of an IDP in physiological conditions with conformational restraints/validation imposed by NMR, SAXS, and smFRET. Using the disordered N-terminal region of the Sic1 protein as a test case (see below), we generated new smFRET and SAXS data to complement previously published NMR data. ^26,27^ To combine these datasets, we used the ENSEMBLE approach (Figure 1), which selects a subset of conformations from a large starting pool of conformations to achieve agreement with experimental data. ^17,28,29^ Our final ensembles of Sic1 are consistent with a diverse set of experimental data suggesting that their properties can be examined with a high degree of confidence. This allowed us to examine relationships between average global polymeric descriptions of Sic1 and higher-moments of their distributions.

**Figure 1:**
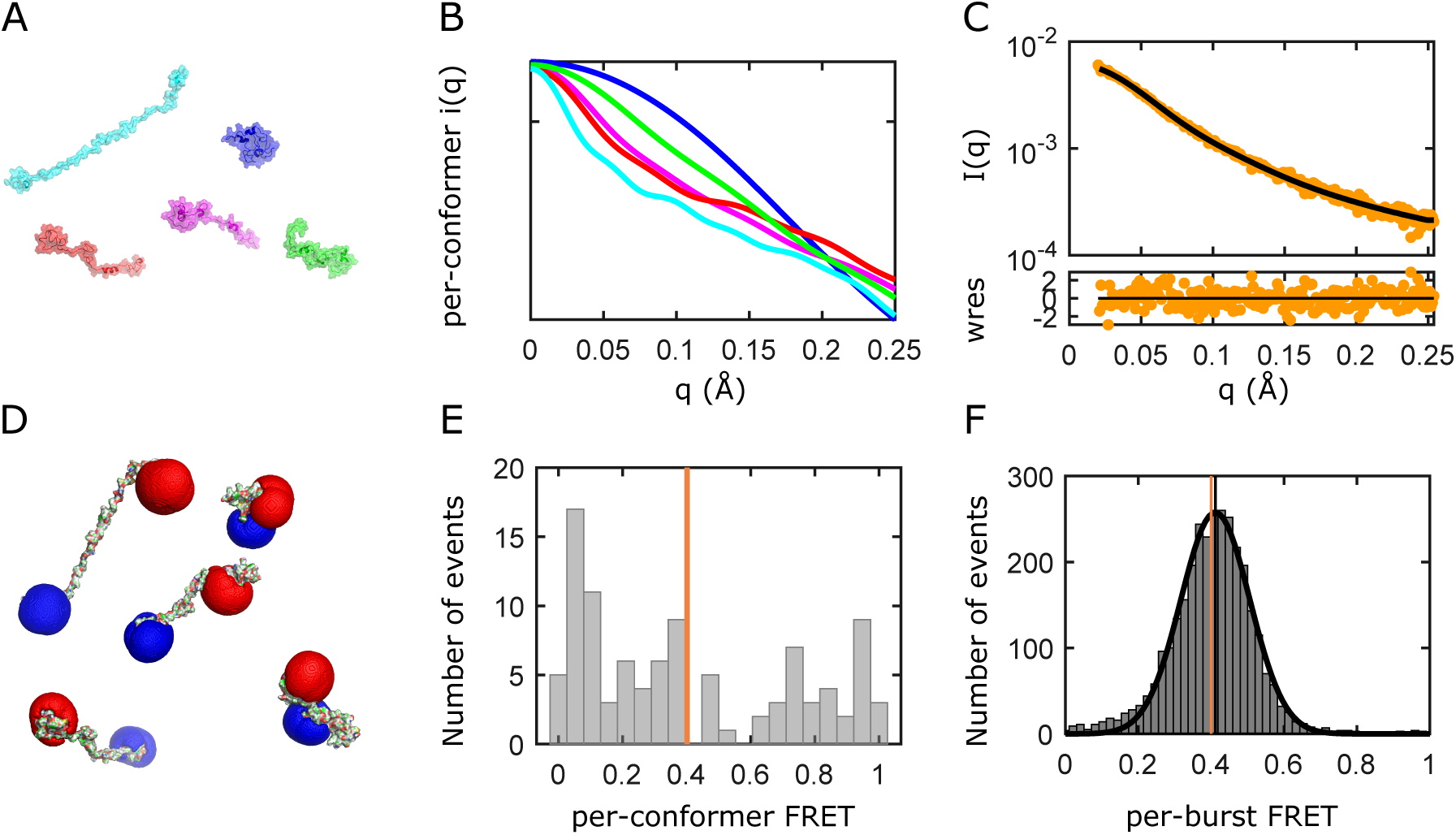
A schematic showing the ENSEMBLE approach for SAXS and smFRET data from an ensemble of structures. (A-B) The SAXS intensity curve of each conformation, *i*(*q*), is back-calculated from the atomic coordinates using CRYSOL. ^30^ (C) The linear average of the CRYSOL-calculated SAXS profiles of individual conformers (black) is compared with the experimental SAXS profile (yellow). (D-E) Per-conformer FRET efficiencies, are calculated assuming a quasi-static distribution of inter-dye distances predicted by accessible volume simulations.^31,32^ (F) The ensemble-averaged transfer efficiency 〈*E*〉_*ens*_ (orange vertical line in E and F) is compared to the mean experimental transfer efficiency 〈*E*〉_*exp*_ (black vertical line).

Achieving our objective of determining Sic1 ensembles consistent with all three datasets also allows us to provide additional insight into the so-called “smFRET and SAXS controversy”. ^33–35^ Previous studies have either *(i)* posited attractive fluorophore interactions in the absence of denaturant, ^22,36^ or *(ii)* have jointly restrained ensemble calculations using both the smFRET and SAXS data.^19,21,37^ The latter approach is based on the recognition that for heteropolymers, deviations from homopolymer chain statistics can cause smFRET and SAXS to be sensitive to different aspects of IDP structure. ^21,38^ For a given IDP and set of labels, both explanations for discrepant inferences are *a priori* plausible and so additional experimental information is needed. Additional experimental information in approach (*ii*) is provided by self-consistent smFRET distance inferences with labels of varying physicochemical properties ^19^ or self-consistent SAXS measurements of samples with and without FRET labels. ^21^ Rather, we provide additional experimental information in the form of NMR restraints, and reserve the smFRET data as an independent validation. Consistency with the smFRET data indicates that globally, perturbative effects of PRE or smFRET labels on the Sic1 ensemble are minimal.

In yeast, the disordered protein Sic1 is eliminated via ubiquitination by the SCF^Cdc4^ ubiquitin ligase and subsequent degradation by the proteasome, allowing initiation of DNA replication. ^39,40^ Sic1 binding to Cdc4, the substrate recognition subunit of the ubiquitin ligase, generally requires phosphorylation of a minimum of any six of the nine Cdc4 phosphodegron (CPD) sites on (full length) Sic1. This effectively sets a high threshold for the level of active G1 CDK required to initiate transition to S-phase. This ultrasensitivity with respect to G1 CDK activity ensures a coordinated onset of DNA synthesis and genomic stability. ^39^ The N-terminal 90 residues of Sic1 (henceforth Sic1) are sufficient for targeting to Cdc4 when highly phosphorylated (henceforth pSic1), making this region a valuable model for structural characterization.^41^ Neither phosphorylation, nor binding to Cdc4 leads to folding of Sic1. ^26,27^ As the binding properties of Sic1 and pSic1 are vastly different, accurate conformational ensembles of Sic1 and pSic1 are central to developing and validating biophysical models of their differential binding. ^42–44^

Surprisingly, previous analysis showed only subtle global changes in Sic1 upon phosphorylation, though only SAXS data was used to restrain the global dimensions. The insensitivity of global dimensions to phosphorylation is surprising given the drastic changes in charge, but is consistent with proposed polyelectostatic models of ultrasensitivty. ^42^ These subtle changes resemble those of another yeast IDP, Ash1, and point to compensatory effects from local and long-range intrachain contacts,^8^ that would be difficult to quantify without an integrative approach. Our integrative modelling, including new smFRET and SAXS data, allows us to examine the details of Sic1 phosphorylation at a previously unattainable level.

## 2 Results

### 2.1 Measurements of *R*_*ee*_ and *R*_*g*_ inferred individually from smFRET or SAXS provide discrepant descriptions of Sic1 and pSic1 conformational ensembles

Figure 2 A-C shows smFRET data measured on the Sic1 FRET construct, which is based on Sic1(1-90) and hereafter called Sic1. This construct was labelled stochastically at its termini with the FRET donor Alexa Fluor 488 and acceptor Alexa Fluor 647 (Förster radius *R*_0_ = 52.2 ± 1.1 Å, details in Supporting Information). The FRET histogram is fit to a Gaussian function to extract the mean transfer efficiency 〈*E*〉_*exp*_, which reports on the inter-dye distance and therefore the end-to-end distance distribution *P* (*r*_*ee*_) as a result of terminal labelling. Multisite phosphorylated Sic1 (pSic1) was generated via overnight incubation with Cyclin A/Cdk2 resulting in predominantly 6- and 7-fold phosphorylated Sic1, with a minor population of 5-fold phosphorylated Sic1 (determined by ESI mass spectrometery). Upon phosphorylation, 〈*E*〉_*exp*_ decreases from 0.42 to 0.36 indicating chain expansion (precision ±0.005, accuracy ±0.02; see Supporting Information).

**Figure 2:**
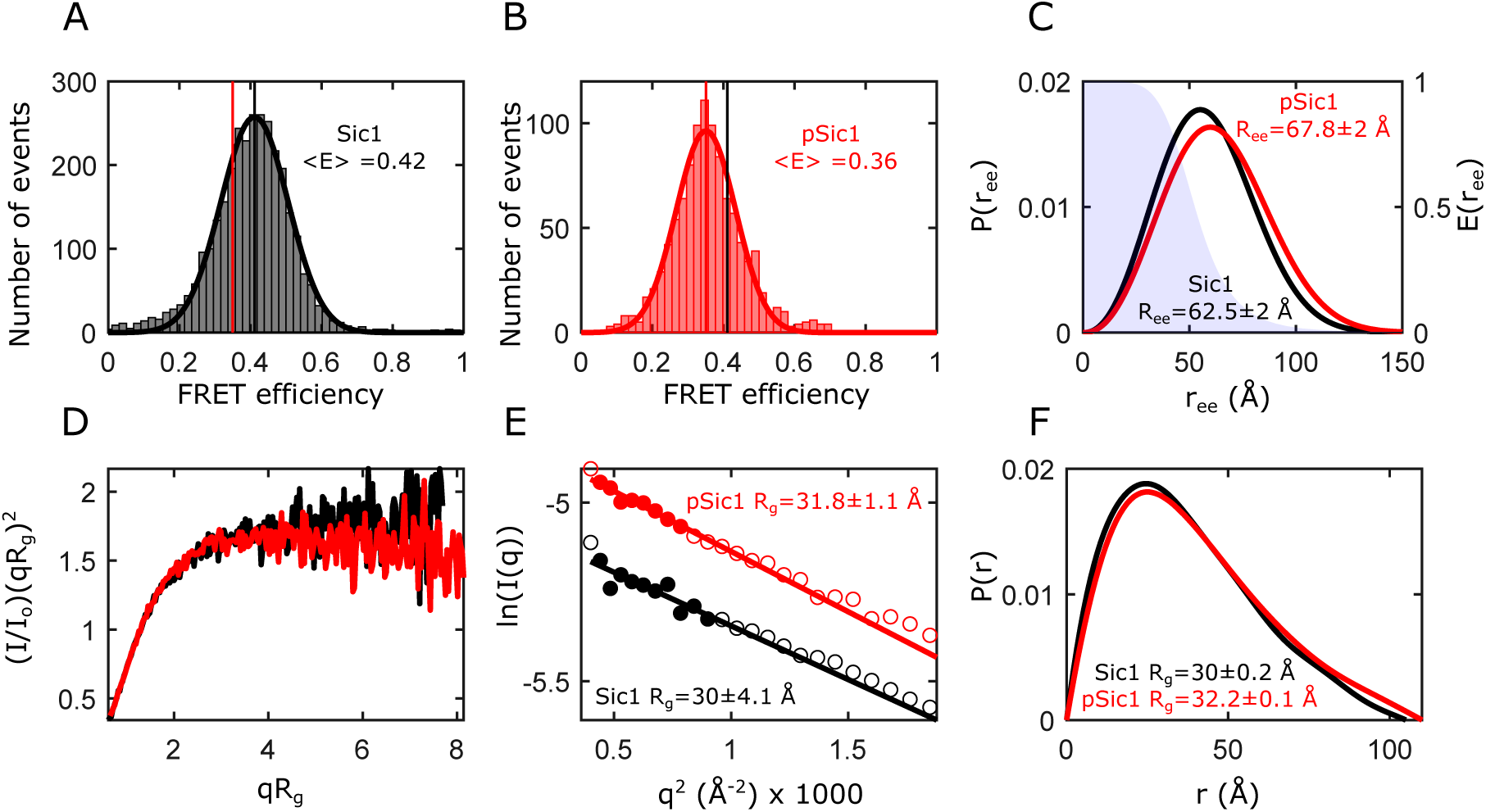
(A-B) smFRET efficiency (*E*) histograms of Sic1 (A) and pSic1 (B) labelled with Alexa Fluor 488 and Alexa Fluor 647 at positions -1C and T90C in TE buffer pH 7.5 150 mM NaCl. (C) Example SAW homopolymer *P* (*r*_*ee*_) distributions (left vertical scale) for Sic1 (black) and pSic1 (red). The shaded underlying region shows the FRET distance dependence function *E*(*r*_*ee*_) (right vertical scale). (D) Dimensionless Kratky plots of Sic1 (black) and pSic1 (red), normalized by initial intensity *I*_0_ and the *R*_*g*_ estimated from the DATGNOM ^45^ fit of the distance distribution function. (E) Guinier plots of Sic1 (black) and pSic1 (red). The solid circles are the data points selected for fitting a restricted range appropriate for IDPs (*q*_max_*R*_*g*_ < 0.9) and the solid lines show the Guinier fits using these data points. (F) The normalized distance distribution function *P* (*r*) estimated by DATGNOM for Sic1 (black) and pSic1 (red).

An estimate of the root-mean-squared end-to-end distance *R*_*ee*_ can be made from 〈*E*〉_*exp*_ by assuming *P* (*r*_*ee*_) is described by a homopolymer model (details in Supporting Information). However, the smFRET data itself does not suggest which (if any) homopolymer model is appropriate for a certain IDP. There is considerable flexibility in the choice of homopolymer model and in how to rescale the root-mean-squared inter-dye distance *R*_*D,A*_ to *R*_*ee*_. This results in a range of *R*_*ee*_, 61-65 Å for Sic1 and 66-72 Å for pSic1, which exceeds other sources of uncertainty (Table S2). If the same polymer modelling is used to analyze Sic1 and pSic1, multisite phosphorylation results in an approximately 10% increase in *R*_*ee*_. However, the smFRET data alone cannot justify this assumption.

To infer the root-mean-squared radius of gyration *R*_*g*_ from *R*_*ee*_ requires an additional assumption about the polymeric nature of system under study, namely the ratio 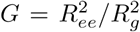, which cannot be determined from the smFRET experiment itself. It has recently been shown that finite-length heteropolymeric chains can take on values of *G* that deviate from the values derived for infinitely long homopolymers in either the *θ*-state (Gaussian chains, *G* = 6) or excluded-volume (EV)-limit (self-avoiding walks, *G* ≈ 6.25). ^21,37,38^ Application of polymer-theoretic values of *G* to the smFRET inferred *R*_*ee*_ results in *R*_*g*_ 24-27 Å for Sic1 and 26-29 Å for pSic1 (Table S3).

Figure 2 D-F shows SAXS data for Sic1 and pSic1. *R*_*g*_ was estimated to be approximately 30 Å for Sic1 and 32 Å for pSic1 using the Guinier approximation, and from the distance distribution function *P* (*r*) obtained using the indirect Fourier transform of the regularized scattering curve (Figure 2 E&F and SI text). Though a model of chain statistics does not need to be specified, these methods are limited in describing IDPs and unfolded proteins. ^13,19^ For example, the expanded and aspherical conformations of IDPs lead to a reduced range of scattering angles in which the Guinier approximation can be applied without systematic error. ^19^ The degree of underestimation of *R*_*g*_ increases as the maximum scattering angle *q*_*max*_ increases, while decreasing *q*_*max*_ reduces the number of points restraining the Guinier fit, which increases the uncertainty in *R*_*g*_ ^19^ (see also, Table S4). In particular, for Sic1, the restricted range (*q*_*max*_*R*_*g*_ < 0.9) which gives similar *R*_*g*_ to analysis of the full SAXS profile (see below) introduces considerable error in *R*_*g*_ (±4 Å).

One solution to these limitations is to model the protein chain explicitly by generating ensembles of conformations. This is epitomized by the Ensemble Optimization Method (EOM) ^46^ and ENSEMBLE method. ^28^ Both approaches select a subset of conformations from an initial pool of conformations, such that the linear average of the CRYSOL-calculated SAXS profiles of individual conformers is in agreement with the full experimental SAXS profile (Figure 1 A-C). However, the techniques differ in their generation of the initial pool of conformations and in the algorithm and cost-function used to minimize the disagreement with experiment (details in the Supporting Information). Despite their differences, both ensemble-based approaches fit the SAXS data equally well, and resulted in nearly identical *R*_*g*_ values, which are similar to the “model-free” estimates (Table S5). As was seen from the smFRET data, multisite phosphorylation results in chain expansion; the SAXS data indicates an approximately 6% increase in *R*_*g*_.

Riback and coworkers have recently introduced another procedure for fitting SAXS data, by pre-generating ensembles of conformations with different properties (specifically, the strength and patterning of inter-residue attractions) and extracting dimensionless “molecular form factors” (MFFs). ^22,36^ The properties of interest are then inferred from the ensemble whose MFF best fits the data. Using the MFFs generated from homopolymer or heteropolymer simulations results in similar *R*_*g*_ to the aforementioned methods (Table S6). In summary, *R*_*g*_ is strongly determined by the SAXS data, such that differences in the construction and refinement of models leads to minor differences in *R*_*g*_.

Analysis of the full SAXS profiles using conformational ensembles shows that the smFRET and SAXS data, analysed individually, provide discrepant inferences of Sic1 and pSic1 global dimensions. Although the various methods calculate ensembles which fit the SAXS data equally well, they have distinct values of *R*_*ee*_, i.e., from 71-81 Å for Sic1 and from 71-87 Å for pSic1 depending on the method used (Tables S5&6). Unlike *R*_*g*_, the SAXS data does not uniquely determine *R*_*ee*_, independent of modelling approach. Taking the ENSEMBLE SAXS-only inferred *R*_*ee*_ as representative, the inferred *R*_*ee*_ = 76.0 ± 2 Å (SEM, 5 replicates) for Sic1 is larger than the largest smFRET inferred *R*_*ee*_ = 65.4 ± 2 Å. Similarly, for homopolymer-based smFRET inferences, the largest Sic1 *R*_*g*_ = 26.8±1.6 Å is still smaller than the SAXS inferred *R*_*g*_ = 30.1±0.4 Å (SEM, 5 replicates) using the ENSEMBLE method.

The benefits of integrative modelling are apparent from the above analysis. Naturally, the accuracy of those aspects of the ensemble not strongly determined by the SAXS data will depend on the initial conformer generation and the optimization/selection algorithms. The wide range of SAXS-inferred *R*_*ee*_ suggests that integrating additional experimental data will improve weakly restrained structural properties, possibly reducing the discrepancy with smFRET. Likewise, 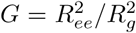 cannot be determined from either dataset individually, and must be assumed *a priori* in smFRET or influenced by assumptions inherent to each SAXS analysis method. It would therefore be desirable to back-calculate a mean FRET efficiency 〈*E*〉_*ens*_ from a structural ensemble that is restrained by SAXS and additional experimental data and to compare 〈*E*〉_*ens*_ and 〈*E*〉_*exp*_ directly. Finally, although the differences in 〈*E*〉_*exp*_ for Sic1 and pSic1 are significant (Δ〈*E*〉_*exp*_ = 0.065 ± 0.007) their *R*_*ee*_ cannot be compared with commensurate precision, since the same homopolymer model may not apply to both.

### 2.2 Ensembles jointly restrained by SAXS and NMR data are consistent with measured FRET efficiencies

We hypothesized that jointly restraining ensembles with non-smFRET internal distance restraints and SAXS data could result in ensembles with back-calculated mean transfer efficiencies, 〈*E*〉_*ens*_, in agreement with the experimental mean transfer efficiency 〈*E*〉_*exp*_. In addition to independently validating the calculated ensemble, this would provide compelling evidence that the smFRET and SAXS data sets are mutually consistent.

To provide non-smFRET information for joint refinement with SAXS data we used previously published NMR data on Sic1. ^26,27^ Briefly, the NMR data consist of ^13^C_*α*_ and ^13^C_*β*_ chemical shifts (CSs) from Sic1 and Paramagnetic Relaxation Enhancement (PRE) data from six single-cysteine Sic1 mutants using a nitroxide spin label (MTSL) coupled to cysteine residues in positions −1, 21, 38, 64, 83, and 90. We used the ENSEMBLE approach to calculate ensembles that are in agreement with the NMR and SAXS data (see Materials and Methods and Supporting Information). We used fluorophore accessible volume (AV) simulations ^31^ to back-calculate the mean transfer efficiency 〈*E*〉_*ens*_ from the sterically accessible space of the dye attached to each conformation via its flexible linker (details in Supporting Information). Briefly, the back-calculated 〈*E*〉_*ens*_ are averages over the accessible inter-dye distances for a particular conformation, as well as averages over all conformations in an ensemble. To determine the proper time-averaging regime, we performed Monte-Carlo simulations of the photon emission process and Brownian motion simulations of dye translational diffusion within the space allowed by sterics and its flexible linker. The slow inter-dye and end-to-end distance dynamics, relative to the donor excited state lifetime, allows 〈*E*〉_*ens*_ to be calculated using the quasi-static averaging approximation.

Table 1 summarizes the agreement of the Sic1 ensembles under various restraint and validation combinations. The agreement of the experimental and back-calculated NMR and SAXS data was quantified using a reduced *χ*^2^ inspired metric. This metric gives an impression of the level of agreement with the various data, though a number of assumptions required for *χ*^2^ statistics are only approximately held. Strictly speaking, reduced *χ*^2^ ∼ 1 indicates a good fit, only if the weighted residuals are standard normally distributed, and the degrees of freedom can be accurately estimated. For the PRE restraints, we have allowed generous error margins (±5 Å) to account for our highly simplified treatment of PRE restraints, and so do not expect standard normally distributed PRE residuals. Similarly, for the CS restraints, the prediction errors derived from training and validation on folded proteins may not accurately predict errors for IDPs. ^47,48^ For SAXS, there are difficult to quantify back-calculation uncertainties from implicit hydration modelling.^49^ For all measurements, the degrees of freedom can be smaller than the number of data points, because of correlations in the data^50^ and the selection of conformers may be considered as free parameters. However, since we likely overestimate errors and degrees of freedom, *χ*^2^ >> 1 indicates disagreement with experiment. The above concerns do not prevent using *χ*^2^ for model comparison. As a structureless null-hypothesis we also include a random coil (RC) ensemble generated with the statistical coil generator TraDES for Sic1. ^51,52^ Residue-by-residue fits to the NMR restraints are shown in Figures S5-7, and fits to the full SAXS profile in Figure S8.

**Table 1:** Agreement of Sic1 *N*_*conf*_ = 500 ensembles with experimental data ^a^

| Restraints | $\chi^2$ PRE | $\chi^2$ $^{13}\text{C}_\alpha$ CS | $\chi^2$ $^{13}\text{C}_\beta$ CS | $\chi^2$ SAXS | $\langle E \rangle_{exp} - \langle E \rangle_{ens}$ |
| --- | --- | --- | --- | --- | --- |
| TraDES RC (none) | 1.51 | 0.514 | 0.518 | 2.03 | 0.12 |
| SAXS | 3.56 | 0.578 | 0.575 | 0.952 | 0.17 |
| PRE | 0.230 | 0.607 | 0.632 | 14.03 | -0.18 |
| SAXS+PRE | 0.252 | 0.511 | 0.462 | 1.01 | 0.02 |
| SAXS+PRE+CS | 0.246 | 0.456 | 0.185 | 0.986 | -0.003 |

<sup>a</sup> $N_{conf} = 500$ ensembles are derived by combining conformations from five independently calculated $N_{conf} = 100$ ensembles. Differences $|\langle E \rangle_{exp} - \langle E \rangle_{ens}| \leq \sqrt{\sigma_{E,exp}^2 + \sigma_{E,ens}^2} \approx 0.02$ indicate no disagreement between back-calculated and experimental mean transfer efficiencies (see Materials and Methods).

The TraDES random coil (RC) ensemble agrees with the CS data, however, the agreement with the PRE, smFRET and SAXS data is poor. Internal distances between specific residues are generally larger in the RC ensemble than are expected from the PRE and smFRET data. The RC ensemble has significant discrepancy in the low-q region, as it underestimates the radius of gyration (Figure S8 and Table S11). When only the SAXS data are used as a restraint, the ensemble reproduces the SAXS curve very well. However, relative to the RC ensemble, the *overall* larger inter-residue distances in the SAXS ensemble further deteriorate the agreement with data reporting on specific inter-residue distances (PRE: Figure S5 and Table 1; smFRET: Table 1 and S9).

When only the PRE data are used as a restraint, the agreement with the PRE data is achieved at the expense of not agreeing with all other observables. This ensemble reproduces specific inter-residue distances encoded by the PRE data, but not the overall distribution of inter-residue distances encoded by the SAXS data. A corollary of the *r*^−6^ PRE weighting is that the PRE ensemble average is dominated by contributions from compact conformations. ^53^ Consistent with this, the PRE-only ensemble is much more compact (*R*_*g*_ ≈ 22 Å) than expected from the SAXS data. Similarly, the transfer efficiency calculated from the ensemble 〈*E*〉_*ens*_ is larger than 〈*E*〉_*exp*_ indicating either too short end-to-end distances overall, or some conformations with strongly underestimated end-to-end distances. Although the absolute value of *χ*^2^ suggests agreement with CSs, the PRE-only ensemble is in worse agreement with the CS data than the TraDES RC or SAXS ensemble.

When the overall distribution of inter-residue distances from SAXS and the specific pattern of interresidue distances from PRE are synthesized in one ensemble model, the transfer efficiency calculated from the ensemble, 〈*E*〉_*ens*_, is in excellent agreement with the experimental transfer efficiency, 〈*E*〉_*exp*_. The fit of the CS data (which were not used as a restraint for this ensemble) are also improved relative to the TraDES RC, the SAXS, and the PRE ensembles. As was previously observed, generating ensembles by satisfying tertiary structure restraints seems to place some restraints on the backbone conformations. ^54^ Finally, we calculated ensembles jointly restrained by SAXS, PRE, and CS data. This improves agreement with CSs, in particular *C*_*β*_ CSs (Figure S7), while the agreement with the other experimental data is within the variation for SAXS+PRE calculations.

### 2.3 Integrative modelling provides a richer description of global dimensions than can be provided by SAXS or smFRET individually

To better understand the implications and advantages of combining multiple datasets we calculated global descriptions of Sic1 and pSic1 conformational ensemble dimensions (radius of gyration *R*_*g*_, end-to-end distance *R*_*ee*_, and hydrodynamic radius *R*_*h*_). Table S11 summarizes the global dimensions of five independently calculated ensembles with 100 conformations each (*N*_*conf*_ = 100).

The radii of gyration, including the implicit solvent layer, of the SAXS+PRE and SAXS+PRE+CS ensembles are ≈ 5% smaller than the SAXS-only estimates. However, no attempt was made to optimize the default solvation parameters in CRYSOL, and small differences in these parameter can result in a 5% to 10% change in *R*_*g*_ for the same set of protein coordinates. ^49^ The radius of gyration calculated directly from the *C*_*α*_ coordinates of the fully restrained ensembles is ≈ 29.5 ± 0.1 Å and ≈ 30.0 ± 0.1 Å for Sic1 and pSic1 respectively (SEM, 5 replicates). As Sic1 and pSic1 have larger than random coil (excluded volume) *R*_*g*_ (i.e., 27.9 ± 0.2 Å; Table S11), we focus on the performance of the self-avoiding walk (SAW) homopolymer models to infer end-to-end distances *R*_*ee*_. The Gaussian chain model has a known tendency to overestimate *R*_*ee*_ when the underlying chain statistics are closer to those of an excluded volume polymer. ^24,38,55^ The end-to-end distance of the fully restrained/validated Sic1 and pSic1 ensembles is ≈ 62 ± 1 Å and ≈ 69 ± 1 Å respectively (SEM, 5 replicates). The SAW homopolymer model inferences of *R*_*ee*_ agree within error, with an average percent error of 1% and −2% for Sic1 and pSic1 respectively.

The above analysis shows that individually, SAXS and smFRET, can accurately infer *R*_*g*_ and *R*_*ee*_ respectively. However, we wish to highlight the advantages of an integrative analysis for Sic1 and pSic1. The global conformational properties of pSic1, as measured by SAXS, are very similar to those of Sic1. This is surprising, given the change in the net charge per residue from ca. 0.12 to −0.01 and −0.03 for 6- and 7-fold phosphorylated Sic1. However, this global insensitivity to phosphorylation state has been observed in a similar yeast IDP, Ash1, ^8^ and is required in the polyelectrostatic model of Sic1 ultrasensitive binding to Cdc4. ^42^ The SAXS ensembles suggest that *R*_*ee*_ is similarly insensitive to multisite phosphorylation (Table S11), while the jointly restrained SAXS+PRE ensembles show an expansion that is confirmed by a direct measurement, smFRET. Similarly, two-dimensional scaling maps (see below) point to heterogeneous changes in internal distances upon phosphorylation that could be observed/validated by future smFRET measurements.

The calculated hydrodynamic radius, *R*_*h*_, was found to be highly similar for all considered ensembles (*R*_*h*_ ≈ 21 − 23 Å). Although we can determine *R*_*h*_ with high precision (variation between replicates is very small < 0.3 Å, Table S11) the accuracy is considerably lower. There are larger margins of error back-calculating a dynamic quantity (*R*_*h*_) from a set of static structures, and in how to properly model solvation effects. For example, calculating *R*_*h*_ using the Kirkwood-Riseman approximation^56^ or using HYDROPRO ^57^ result in *R*_*h*_ values that differ by ca. 20%. Thus, while it is encouraging that ensemble *R*_*h*_ are close to experimental values determined by NMR ^27^ (*R*_*h*_ = 21.5 ± 1.1 Å for Sic1 and *R*_*h*_ = 19.4 ± 1.6 Å for pSic1) and by FCS ^58^ (*R*_*h*_ = 22 ± 2 Å for Sic1) it is premature to consider this a validation of the ensemble, especially given the insensitivity of *R*_*h*_ to different restraint combinations (see Table S11).

### 2.4 Analysis of the conformational behaviour of calculated ensembles beyond global dimensions

We next sought to determine descriptions of the calculated conformational ensembles which go beyond global dimensions and would facilitate comparison with polymer theory reference states, and with IDPs and unfolded states of varying sequence and chain length, *n*. To this end, we used the fact that many aspects of homopolymer behaviour become universal, or independent of monomer identity, in the long chain (as *n → ∞*) limit. ^59^ This allowed us to clearly identify ways in which ensembles deviate from homopolymer behaviour (Table 2).

**Table 2:**
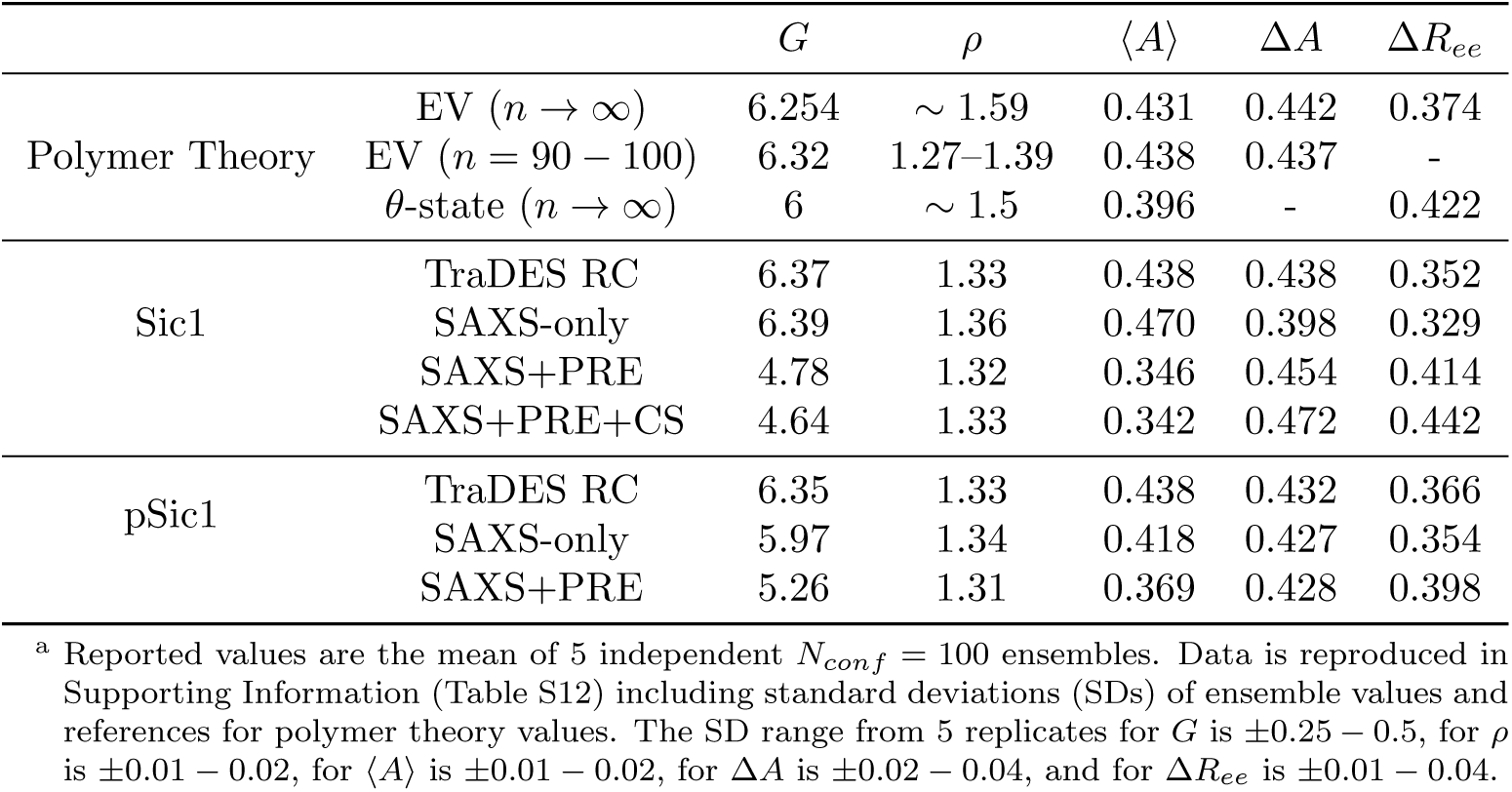
Nominally universal polymer properties of the calculated ensembles^a^

| | | $G$ | $\rho$ | $\langle A \rangle$ | $\Delta A$ | $\Delta R_{ee}$ |
| --- | --- | --- | --- | --- | --- | --- |
| Polymer Theory | EV ($n \rightarrow \infty$) | 6.254 | $\sim 1.59$ | 0.431 | 0.442 | 0.374 |
| | EV ($n = 90 - 100$) | 6.32 | 1.27–1.39 | 0.438 | 0.437 | - |
| | θ-state ($n \rightarrow \infty$) | 6 | $\sim 1.5$ | 0.396 | - | 0.422 |
| Sic1 | TraDES RC | 6.37 | 1.33 | 0.438 | 0.438 | 0.352 |
|  | SAXS-only | 6.39 | 1.36 | 0.470 | 0.398 | 0.329 |
|  | SAXS+PRE | 4.78 | 1.32 | 0.346 | 0.454 | 0.414 |
|  | SAXS+PRE+CS | 4.64 | 1.33 | 0.342 | 0.472 | 0.442 |
| pSic1 | TraDES RC | 6.35 | 1.33 | 0.438 | 0.432 | 0.366 |
|  | SAXS-only | 5.97 | 1.34 | 0.418 | 0.427 | 0.354 |
|  | SAXS+PRE | 5.26 | 1.31 | 0.369 | 0.428 | 0.398 |

<sup>a</sup> Reported values are the mean of 5 independent $N_{conf} = 100$ ensembles. Data is reproduced in Supporting Information (Table S12) including standard deviations (SDs) of ensemble values and references for polymer theory values. The SD range from 5 replicates for $G$ is $\pm 0.25 - 0.5$ , for $\rho$ is $\pm 0.01 - 0.02$ , for $\langle A \rangle$ is $\pm 0.01 - 0.02$ , for $\Delta A$ is $\pm 0.02 - 0.04$ , and for $\Delta R_{ee}$ is $\pm 0.01 - 0.04$ .

For very long homopolymer chains, the scaling exponent *ν* tends to one of only three possible limits (1/3, 1/2, 0.588), describing the poor-solvent, *θ*-state, and excluded volume (EV)-limit respectively. Homopolymers in these limits have well-defined universal values for the size ratios 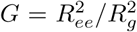 and *ρ* = *R*_*g*_*/R*_*h*_, the overall shape of the ensemble, as characterized by the average asphericity 〈*A*〉 (*A* ∼ 0 for a sphere and *A* ∼ 1 for a rod), the relative variance in the end-to-end distance distribution 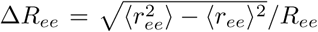, and the relative variance in the distribution of the shape of individual conformations 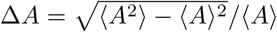. Table 2 summarizes the universal values expected for homopolymers in the *θ*-state or the EV-limit, in the case of very long chains (EV and *θ*-state *n → ∞*) and for chains with similar length to Sic1 (EV *n* = 90−100). The TraDES random coil, though not a homopolymer, is constructed with only excluded volume long-range interactions, and so is expected to have behaviour consistent with polymer theory predictions for an EV-limit polymers of similar chain-length (EV *n* = 90 − 100 Table 2).

The values of *G* for the Sic1 and pSic1 random coil and SAXS-only ensembles are indistinguishable from the expected value for a homopolymer in the EV-limit (*G* ≈ 6.3). In contrast, ensembles jointly restrained by SAXS and PRE have *G* outside the range *G*_*θ*_ = 6 *≤ G ≤ G*_*EV*_ ≈ 6.3 despite having apparent scaling exponents between the *θ*-state and EV-limits (see below). For Sic1 *G* = 4.7 ± 0.1 and for pSic1 *G* = 5.3 ± 0.1 (SEM, 5 replicates). The ratio *ρ*, on the other hand, is not sensitive to deviations from homopolymer statistics at long sequence separations. The value of *ρ* remains ∼1.3 for all ensembles, despite large changes in *R*_*ee*_ and *G*. The calculated *ρ* are consistent with the range of polymer-theoretic values for a finite length EV homopolymer (EV *n* = 90 − 100 Table 2).

The Sic1 and pSic1 RC and SAXS-only ensembles, have an average asphericity 〈*A*〉 very close to the polymer-theoretic value for a homopolymer in the EV-limit. Although individual conformations are not necessarily spherical, SAXS+PRE ensembles of both Sic1 and pSic1 are on average more spherical, with significantly lower 〈*A*〉, despite their larger-than-RC *R*_*g*_. Similar to *G*, the values of 〈*A*〉 for the SAXS+PRE ensembles (0.346 ± 0.005 for Sic1 and 0.369 ± 0.005 for pSic1, SEM 5 replicates) are outside of the *θ*-state and EV-limit.

The relative variance in the end-to-end distance distribution, Δ*R*_*ee*_ is close to the EV-limit value 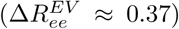 for the random coil and SAXS-only restrained ensembles. In contrast, Sic1 and pSic1 SAXS+PRE restrained ensembles have Δ*R*_*ee*_ which are more consistent with the *θ*-state value. Although Sic1 and pSic1 *R*_*ee*_ are more compact than the RC, they exhibit larger relative variations in the end-to-end distances of their conformations. All ensembles have a relative variance in the distribution of shapes, Δ*A*, similar to that of an EV-limit homopolymer. The broadness of the shape distribution stresses the fact that despite being, as an ensemble of conformations, more spherical than an EV polymer, the Sic1 ensembles contains individual conformations with a large distribution of shapes.

### 2.5 Internal scaling profiles and apparent scaling exponents

Recently, the focus of the smFRET and SAXS debate has moved from inferring *R*_*g*_ to inferring apparent scaling exponents. ^22,37^ To extract further insights regarding the effects of combining multiple solution data types on the statistics of internal distances in the ensembles, we calculated internal scaling profiles (ISPs, Figure 3). ISPs quantify the mean internal distances 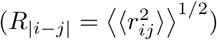 between all pairs of residues that are |*i* − *j*| residues apart in the linear amino acid sequence (see Materials and Methods). The dependence of *R*_|*i*−*j*|_ on sequence separation |*i* − *j*| is often quantified by fitting to the power-law relation:

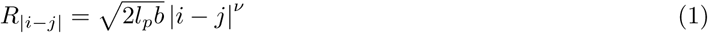

where *b* = 3.8 Å is the distance between bonded C_*α*_ atoms and *l*_*p*_ ≈ 4 Å is the persistence length. This persistence length was found to be applicable to a broad range of denatured and disordered states. ^5,21,60^ Scaling laws are derived for homopolymers in the infinitely-long-chain limit. For a finite-length heteropolymer we measure merely an apparent scaling exponent *v*_*app*_, however we drop the subscript to aid the clarity of the text.

**Figure 3:**
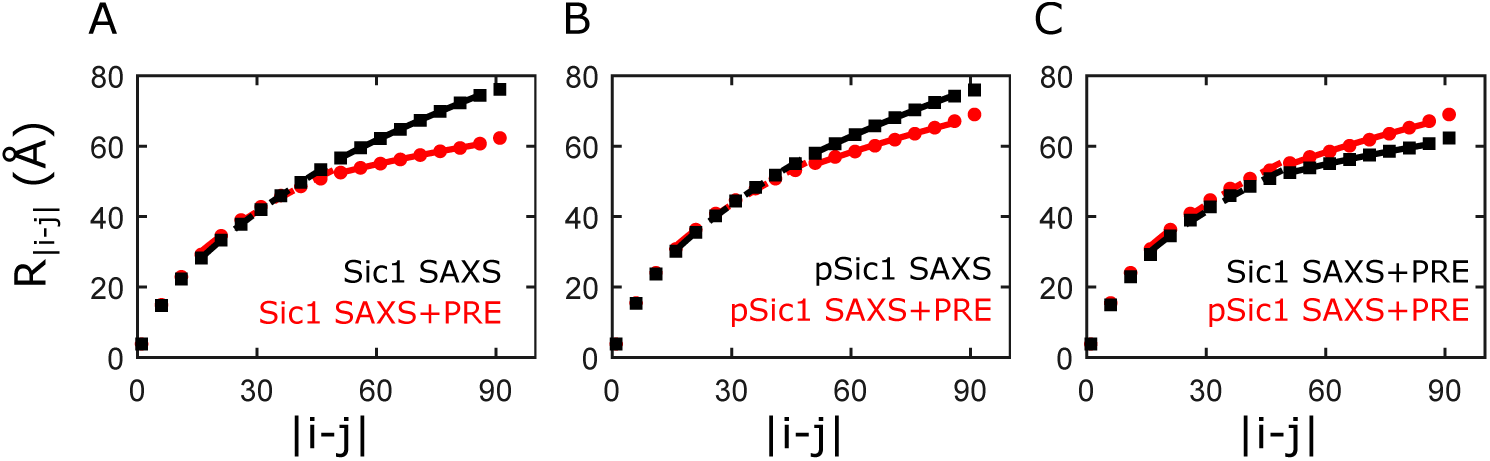
Internal scaling profiles calculated from 5 *N*_*conf*_ = 100 ensembles. (A) Sic1 SAXS+PRE ensembles (red circles) and Sic1 SAXS-only ensembles (black squares). (B) pSic1 SAXS+PRE ensembles (red circles) and pSic1 SAXS-only ensembles (black squares). (C) pSic1 (red circles) and Sic1 (black squares) SAXS+PRE ensembles. For all panels, fits are shown for to intermediate (dashed) and long (solid) sequence separations. For visualization, every fifth data point is shown.

ISPs highlight important differences between ensembles. If the majority of internal distances are similar in two ensembles, their *R* values will be similar, as 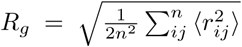. ^21^ However, if their spatial separations start to diverge at long sequence separations, the ensembles will have dissimilar *R*_*ee*_ and 〈*E*〉_*exp*_, when terminally labelled. This decoupling of *R*_*ee*_ from *R*_*g*_ is illustrated by Figure 3A which shows the scaling of the SAXS-only and SAXS+PRE Sic1 ensembles, which have similar *R*_*g*_, but only the SAXS+PRE ensemble is consistent with the smFRET data.

We quantify the change in scaling behaviour at long sequence separations (*v*_*long*_, 51 < |*i* − *j*| *≤ n*_*res*_ − 5) relative to intermediate sequence separations (*v*_*int*_, 15 *≤* |*i* − *j*| *≤* 51) by calculating Δ*v*_*ends*_ = *v*_*long*_ − *v*_*int*_ (5 replicates with *N*_*conf*_ = 100, Table 3). For homopolymers in the long-chain limit we expect Δ*v*_*ends*_ = 0; though finite-length, the Sic1 and pSic1 random coil ensembles have Δ*v*_*ends*_ ≈ 0 (Δ*v*_*ends*_ = −0.06 ± 0.03, SEM 5 replicates). For Sic1, both the SAXS and SAXS+PRE ensembles show Δ*v*_*ends*_ < 0, though the deviation from homopolymer statistics is stronger in the SAXS+PRE ensembles (Δ*v*_*ends*_ = −0.08 ± 0.01 and Δ*v*_*ends*_ = −0.25 ± 0.04 respectively, SEM 5 replicates). Internal distances in the Sic1 SAXS+PRE ensemble follow marginally good-solvent scaling at intermediate sequence separations, and transition to poor solvent scaling at larger sequence separations. Expansion of Sic1 upon phosphorylation has been attributed to transient tertiary contacts involving non-phosphorylated CPDs that are lost or weakened upon phosphorylation. ^26^ Consequently, while pSic1 SAXS ensembles do not identify deviations from homopolymer statistics (Δ*v*_*ends*_ = −0.09 ± 0.05), pSic1 SAXS+PRE ensembles identify smaller deviations than those observed for Sic1 (Δ*v*_*ends*_ = −0.13 ± 0.03).

**Table 3:** Fitting results for the TraDES RC ensemble, SAXS-only ensemble, and SAXS+PRE ensembles ISPs ^a^

|  | Sic1 |  |  | pSic1 |  |
| --- | --- | --- | --- | --- | --- |
|  | TraDES RC<sup>b</sup> | SAXS-only | SAXS+PRE | SAXS-only | SAXS+PRE |
| $\nu$ (fixed $l_p = 4$ Å) | 0.570 | 0.589 | 0.567 | 0.596 | 0.583 |
| $\nu_{int}$ | 0.566 | 0.601 | 0.524 | 0.569 | 0.517 |
| $\nu_{long}$ | 0.51 | 0.52 | 0.28 | 0.47 | 0.38 |
| $\Delta\nu_{ends}$ | -0.06 (0.03) | -0.08 (0.01) | -0.25 (0.04) | -0.09 (0.05) | -0.13 (0.03) |

<sup>a</sup> Table results are the mean results from fitting 5 $N_{conf} = 100$ ensembles. SEM for $\nu$ and $\nu_{int}$ is $\approx 0.005$ and 0.03 for $\nu_{long}$ . SEM for $\Delta\nu_{ends}$ shown in parenthesis. See Materials and Methods for additional details.
<sup>b</sup> Sic1 TraDES RC and pSic1 TraDES RC result in nearly identical fits.

Further, we compared scaling behaviour determined by our integrative approach to recently published methods based only on SAXS data using “molecular form factors” (MFFs) ^22,36^ (Table S6), or only on smFRET data using the SAW-*v* method ^55^ (Table S9). For Sic1, but not pSic1, there is agreement between global scaling determined from the SAXS data and the global scaling determined from the SAXS+PRE ensemble scaling profiles. Due to the terminal labelling positions and because Δ*v*_*ends*_ < 0, *v* inferred from smFRET is less than *v* inferred from SAXS. However, neither approach using a single data type fully captures the heteropolymeric behaviour of Sic1 and pSic1.

### 2.6 Two dimensional scaling maps reveal regional biases for expansion and compaction

To better describe the heteropolymeric nature of Sic1, a normalized two-dimensional (2D) scaling map was constructed (Figure 4). In the first step, the ensemble-averaged distances between the C_*α*_ atoms of every unique pair of residues in the sequence is calculated for the experimentally-restrained ensemble (〈*r*_*ij*_〉_*ens*_), and for the respective TraDES random coil (RC) ensemble (〈*r*_*ij*_〉_RC_). Experimentally-restrained distances are normalized by the RC distances and displayed as a 2D scaling map.

**Figure 4:**
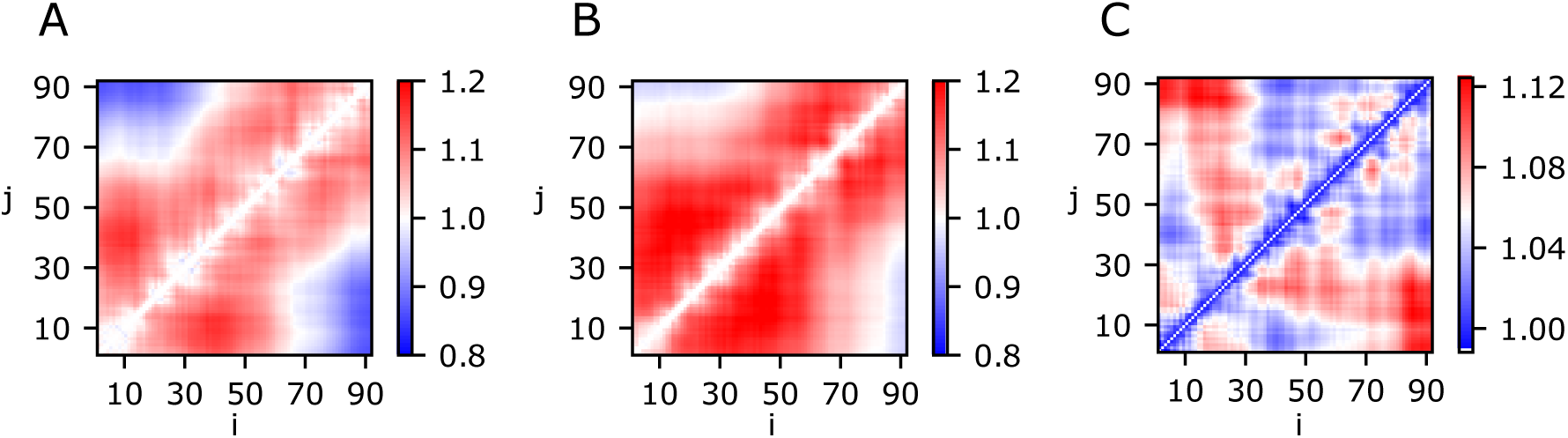
(A) Sic1 2D scaling map *α*_*ij*_ = 〈*r*_*ij*_〉_*ens*_*/*〈*r*_*ij*_〉_*RC*_ using the Sic1 (SAXS+PRE) *N*_*conf*_ = 500 and the Sic1 *N*_*conf*_ = 500 TraDES RC ensemble. (B) pSic1 2D scaling map *α*_*ij*_ = 〈*r*_*ij*_〉_*ens*_*/*〈*r*_*ij*_〉_*RC*_ using the pSic1 (SAXS+PRE) *N*_*conf*_ = 500 and the pSic1 *N*_*conf*_ = 500 TraDES RC ensemble. (C) pSic1 normalized by Sic1 dimensions.

The normalized 2D scaling map for Sic1 (Figure 4A) displays regional biases for expansion (*α*_*ij*_ > 1) and compaction (*α*_*ij*_ < 1). Short internal distances |*i* − *j*| ⪅ 45 show expansion relative to the RC, while |*i* − *j*| ⪆ 60 show compaction. The expansion, however, is heterogeneous. For example, the ∼ 40 residue N-terminal region is more expanded than the ∼ 40 residue C-terminal region. Similar distinctions between the RC and pSic1 ensembles were observed (Figure 4B).

To compare Sic1 and pSic1 ensembles, the pSic1 ensemble was normalized by the Sic1 ensemble, (Figure 4C). This map describes the heterogeneous modulation of Sic1 dimensions upon multisite phosphorylation. Expansion is clustered around CPD sites, particularly those of the C-terminus and in the vicinity of Y14, previously implicated in tertiary interactions with CPDs^27^ (see below).

### 2.7 Y14A mutation and phosphorylation disrupt tertiary contacts in Sic1

We next sought to determine whether specific long-range interactions leading to compact end-to-end distances in Sic1 and pSic1 could be identified and disrupted. PRE effects link CPDs with Y14 and ^15^N relaxation experiments on Sic1 identified maxima in the R_2_ rates near Y14. ^27^ Furthermore, the substitution Y14A led to an expansion in *R*_*h*_ of ∼20% in pSic1. ^27^ We hypothesized that if Y14 engages in specific pi-pi and cation-pi interactions throughout the chain, then removing its pi-character by mutation to alanine will disrupt these interactions, leading to larger *R*_*ee*_ and lower 〈*E*〉_*exp*_.

We performed smFRET experiments for the Y14A mutants of Sic1 and pSic1 (Figure 5 and Table S9). Y14A mutation decreases Sic1 〈*E*〉_*exp*_ by approximately 7% (ca. 0.42 to 0.40, a small but reproducible shift). Phosphorylation of the Y14A mutant decreases its 〈*E*〉_*exp*_ by approximately 16% (ca. 0.40 to 0.33). At this time we cannot rule out that the observed FRET changes may be due (in part) to a different phosphorylation pattern for the Y14A mutant. However, these experiments suggest that the pi-group of Y14 participates in long-range contacts which maintain more compact *R*_*ee*_ in Sic1 and pSic1 than would be expected for a homopolymer with similar *R*_*g*_. These contacts are likely key for the globally compact conformations required in the polyelectrostatic model of pSic1:Cdc4 binding. ^42^ This demonstrates how smFRET can be used to test structural hypotheses generated from integrative modelling.

**Figure 5:**
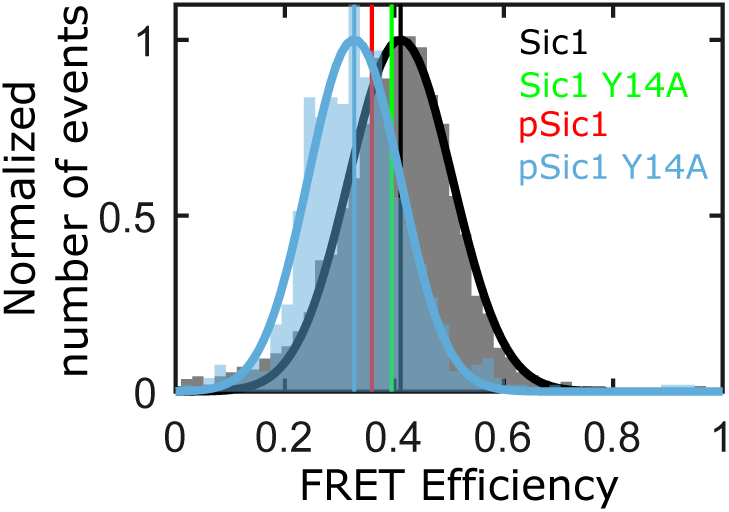
Y14A mutation and phosphorylation results in a shift to lower 〈*E*〉_*exp*_ (more expanded conformations). Each histogram is normalized so that each Gaussian fit has a maximum of one.

**Figure 6:**
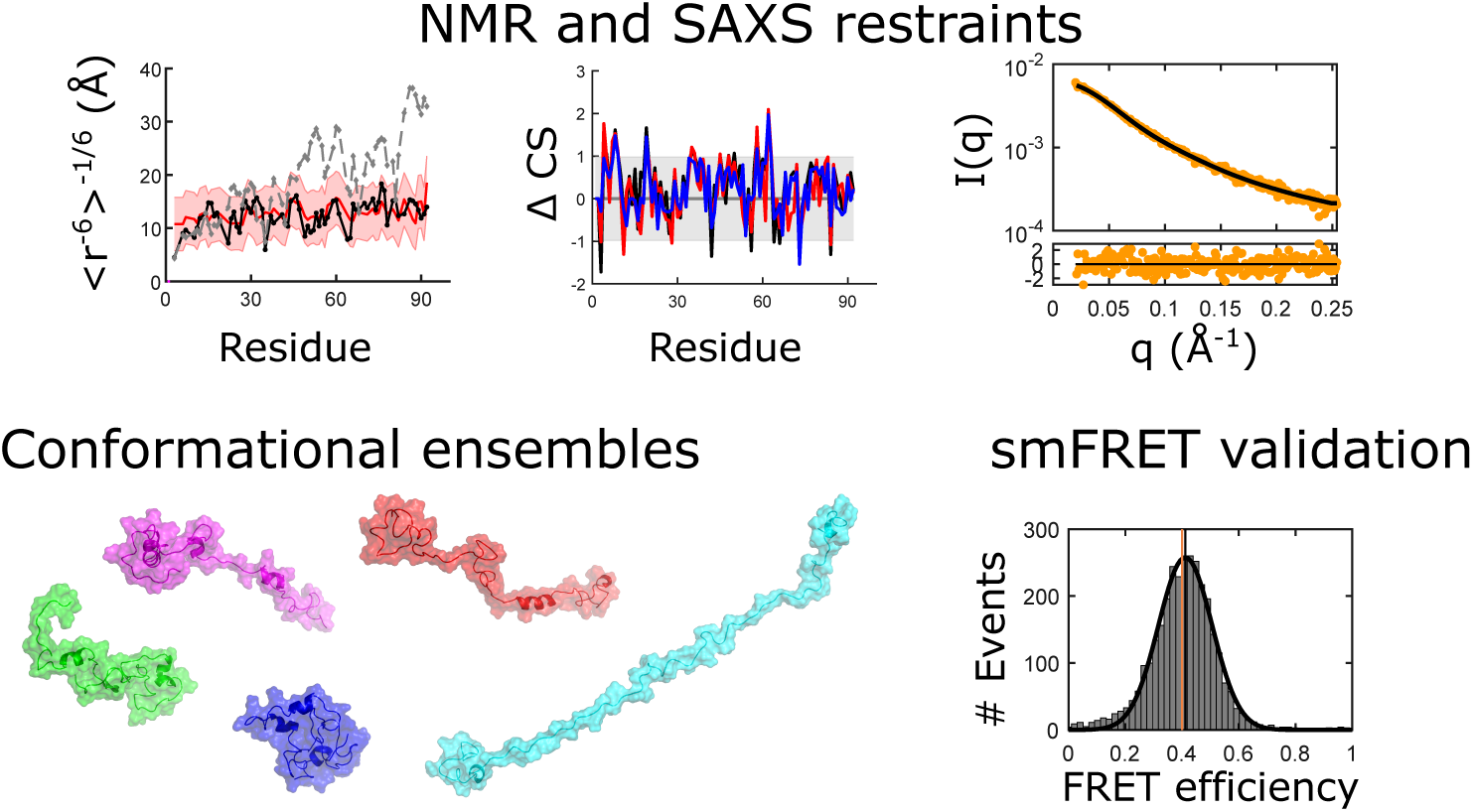
For Table of Contents only

## 3 Discussion

We generated SAXS and smFRET data on Sic1 and pSic1, and resolved their apparently discrepant inferences by joint refinement of the SAXS data with PRE data. The ensembles restrained by SAXS and PRE data are, in addition, consistent with the smFRET data, chemical shift data, and hydrodynamic data (PFG-NMR and FCS). We used smFRET transfer efficiencies directly as validation, rather than using derived distances from the data via polymer theory assumptions. Our final ensembles of Sic1 and pSic1 can be examined with a high degree of confidence given their agreement with a diverse set experimental data acting as both restraints and validation. This was important since the changes in Sic1 upon phosphorylation are quite subtle.

The picture that emerges when the entirety of the experimental data on Sic1 and pSic1 is considered, is that their conformational ensembles cannot be described by statistics derived for infinitely long homopolymers. Although this is unsurprising, given that Sic1 and pSic1 are finite-length heteropolymers, ensembles restrained only by the SAXS data are congruent with the set of homopolymer descriptions and scaling relationships for excluded volume homopolymers. Neither the SAXS nor smFRET data, individually, suggest deviations from homopolymer statistics. Our results therefore provide a strong impetus for integrative modelling approaches over homopolymer approaches whenever multiple data types exist.

We emphasize that the SAXS+PRE ensembles were not constructed by reweighting or selecting ensembles specifically to achieve agreement with 〈*E*〉_*exp*_. In our approach, it was not guaranteed *a priori* that 〈*E*〉_*ens*_ would match 〈*E*〉_*exp*_, especially if either the introduction of PRE spin labels or smFRET fluorophores had perturbed the IDP ensemble. If negative values of Δ*v*_*ends*_ are common for IDPs and for unfolded proteins under refolding conditions, smFRET on terminally labelled samples will infer smaller *v* than would SAXS. When experiments are analysed individually, Δ*v*_*ends*_ < 0 is consistent with both fluorophore-driven interactions and heteropolymer effects. In both cases, Δ*v*_*ends*_ would approach zero in high concentrations of denaturant, which would disrupt both spurious fluorophore interactions and native long-range tertiary interactions. ^21,22^ Deciding between fluorophore-interactions and heteropolymer effects requires an integrative modelling approach. An additional consistency check is to measure 〈*E*〉_*exp*_ for a different pair of dyes with different physicochemical properties. In a previous publication ^58^ we measured 〈*E*〉_*exp*_ for Sic1 using smaller, less charged, and more hydrophobic dyes (TMR and Atto647N). We re-calculated the expected 〈*E*〉_*ens*_ for the *N*_*conf*_ = 500 Sic1 SAXS+PRE ensemble with TMR and Atto647N accessible volumes (Table S7) and the measured Förster radius *R*_0_ = 60 ± 2 Å for this pair. The resulting 〈*E*〉_*ens*_ = 0.51 ± 0.02 agrees with the measured 〈*E*〉_*exp*_ = 0.47 ± 0.02 (Figure 7D of Ref. ^58^).

### 3.1 Conformation-to-function relationships

For soluble post-translationally modified IDPs, approximately good-solvent scaling may be unsurprising. The balance between chain-chain and chain-solvent interactions is a driving force for aggregation ^61^ and phase separation, ^62,63^ and polymer theory predicts that proteins with overall good-solvent scaling in native-like conditions should remain soluble. At short-to-intermediate sequence separations, good-solvent scaling provides read/write access of substrate motifs to modifying enzymes (e.g., phosphorylation and ubiquitination for Sic1).

Good-solvent scaling also confers advantages to dynamic complexes, as internal friction increases with increasing chain compaction. ^64^ Low internal friction and fast chain reconfigurations provides more opportunities for unbound Cdc4 phosphodegrons (CPDs) to (re)bind before pSic1 diffuses out of proximity of Cdc4. ^43,44,65^ In the polyelectrostatic model, fast reconfiguration dynamics facilitates pSic1’s dynamic interactions with Cdc4 through electrostatic averaging effects. ^27,42^

The crossover to poor-solvent scaling at long sequence separations implies that unbound CPDs that are sequence-distant from a bound CPD are on average closer to the WD40 binding pocket than they would be for an EV-chain. This effectively decreases the solvent screening of electrostatic interactions and is predicted to lead to sharp transitions in the fraction bound with respect to the number of phosphorylations. ^42^ Increasing the effective concentration of CPDs in the vicinity of the binding pocket may also increase the probability for any CPD to rebind before diffusive exit.

Large amplitude fluctuations in the shape (Δ*A*) and size (Δ*R*_*ee*_) of Sic1, effectively and rapidly sampling many different conformations, could allow CPDs in Sic1 to rapidly sample either the primary or secondary WD40 binding pocket. These fluctuations could facilitate electrostatic averaging, permitting a mean-field treatment as assumed in the polyelectrostatic model. ^42^

## 4 Conclusions

Our work provides a description of the conformational ensembles of Sic1 and pSic1 which is consistent with experimental data reporting on a wide range of spatial and sequence separation scales, and with biophysical models for Sic1 function. Our results show that there are clear advantages of combining multiple datasets and that quantitative polymer-physics-based characterization of experimentally-restrained ensembles allows the description and classification of IDPs as heteropolymers. The chain length independence of many of these properties facilitates comparison between different IDPs and unfolded states of proteins.

Our results suggest that for Sic1 and our dye pair, discrepant inferences between SAXS and smFRET cannot *a priori* be assumed to arise from “fluorophore-interactions.” The impact of the fluorophores (or spin-labels) will of course depend on the physicochemical properties of the specific IDP sequence and the fluorophores (or spin-labels) used. Robustness to perturbation (e.g., labels or phosphorylation) may be built into Sic1’s sequence via its patterning of charged and proline residues. ^8^ Further understanding of the discriminatory power of FRET, and the utility of different restraint types for characterizing types of structure in IDPs, will come from recently developed Bayesian procedures ^66,67^. In this regard, an integrative use of multiple experiments probing disparate scales, computational modelling, and polymer physics, will provide valuable insights into IDPs/unfolded states and their biological functions.

## 5 Materials and Methods

### 5.1 Sic1 samples

The Sic1 N-terminal region (1-90, henceforth Sic1) was expressed recombinantly as a Glutatione S-transferase (GST) fusion protein in *Escherichia coli* BL21 (DE3) codon plus cells and purified using glutathione-Sepharose affinity chromatography and cation-exchange chromatography. The correct molecular mass of the purified protein was verified by electrospray ionization mass spectrometry (ESI-MS). A double cysteine variant of Sic1 (−1C-T90C) for smFRET experiments was generated via site directed mutagenesis from a single-cysteine mutant produced previously for PRE measurements. ^26,27^ This construct was purified as above and the correct molecular mass of the purified protein was verified by ESI-MS. A Y14A mutant Sic1 (−1C-T90C-Y14A) was generated via site directed mutagenesis from the aforementioned double-cysteine mutant and was expressed, purified, and characterized using the same protocol. Phosphorylated samples were prepared by treatment of Sic1 with Cyclin A/Cdk2 (prepared according to Huang et al., ^68^)at a kinase:Sic1 ratio of 1:100 in the presence of 50 fold excess of ATP and 2.5 mM MgCl_2_ overnight at 30 *°*C. The yield of phosphorylation reaction was determined by ESI-MS. Under these conditions the dominant species are 6- and 7-fold phosphorylated Sic1 (10195 Da and 10274 Da respectively) with a small fraction of 5-fold phosphorylated Sic1. After phosphorylation, the samples were buffer exchanged into PBS buffer pH 7.4 with 3 M GdmCl to prevent aggregation, denature kinase, and denature any phosphatases which may have inadvertently entered the solution. The samples were kept on ice in 4*°*C and measured within 24 hours.

The Sic1 smFRET construct was labelled stochastically with Alexa Fluor 488 *C*_5_ Maleimide (ThermoFisher Scientific, Invitrogen, A10254) and Alexa Fluor 647 *C*_2_ Maleimide (ThermoFisher Scientific, Invitrogen, A20347). After labelling with Alexa Fluor 647, cation-exchange chromatography was used to separate species with a single acceptor label, from doubly acceptor labelled and unlabelled species. The single-labelled species sample was then labelled with Alexa Fluor 488 and cation-exchange chromatography was used to separate doubly heterolabelled from acceptor only species. The correct mass of the doubly labelled sample was confirmed by mass spectrometry. The final FRET labelled sample was concentrated and buffer exchanged into PBS buffer pH 7.4 with 3 M GdmCl, 2 mM DTT and stored at −80 °C. Additional details regarding protein expression, purification and labelling are available in the Supporting Information.

### 5.2 Single-molecule fluorescence

Single-molecule fluorescence experiments were performed on a custom-built multiparameter confocal microscope with microsecond alternating laser excitation. This instrumentation allows the simultaneous detection of the intensity, anisotropy, lifetime, and spectral properties of individual molecules and for the selection of fluorescence bursts in which both dyes are present and photophysically active. The acquired data were subjected to multiparameter fluorescence analysis ^69,70^ and ALEX filtering. ^71^ The burst search was performed using an All Photon Burst Search (APBS) ^72,73^ with *M* = 10, *T* = 500 *µ*s and *L* = 50. Transfer efficiencies were determined burst-wise and corrected for differences in the quantum yields of the dyes and detection efficiencies, as described in further detail in the Supporting Information.

Immediately prior to measurement samples were diluted to ∼50 pM in either (*i*) PBS buffer: 10 mM sodium phosphate and 140 mM NaCl pH 7.0, 1 mM EDTA (to replicate NMR measurement buffer of Ref ^26^) or (*ii*) Tris buffer: 50 mM Tris and 150 mM NaCl, pH 7.5. (to replicate SAXS measurement buffer). No difference in 〈*E*〉_*exp*_ was detected when comparing buffer conditions and results are shown for Tris buffer conditions. Dilution of the smFRET samples from stock concentration in 3M GdmCl to single-molecule concentration results in approximately 60 nM residual concentration of GdmCl. Additionally, the SAXS measurements include 5 mM DTT, and 2 mM TCEP to scavenge radicals and prevent radiation damage but which are detrimental to fluorophore performance; while the smFRET measurements use 143 mM 2-mercaptoethanol (BME, 1:100 v/v dilution) and 5 mM 2-mercaptoethylamine (MEA) for photoprotection and increased brightness. The smFRET samples also contain 0.001 % Tween 20 for surface passivation.

The Förster radius *R*_0_ was calculated assuming a relative dipole orientation factor *κ*^2^ = 2*/*3 and the refractive index of water *n* = 1.33. The assumption of *κ*^2^ = 2*/*3 is supported by subpopulation-specific steady-state anisotropies for the donor in the presence of the acceptor (Table S1). The overlap integral *J* was measured for each sample and found not to change upon phosphorylation or Y14A mutation. The minimal variation in donor-only lifetimes *τ*_*D*0_ suggested minimal variation in the donor-quantum yield *ϕ*_*D*_. *R*_0_ was therefore calculated to be *R*_0_ = 52.2 ± 1.1 Å for all samples, and variation between samples within this uncertainty.

We estimate the precision for 〈*E* 〉_*exp*_ to be ca. 0.005 (for measurements performed on the same day, with approximately equal sample dependent calibration factors). We estimate the accuracy of 〈*E*〉_*exp*_, *σ*_*E,exp*_, to be ca. 0.02 (due to uncertainty in the instrumental and sample dependent calibration factors). Further details about the instrumentation, photoprotection, laser excitations, burst detection, filtering and multiparameter fluorescence analysis can be found in the Supporting Information.

### 5.3 Small-angle X-ray scattering

Small angle X-ray scattering data were collected at beamline 12-ID-B at the Argonne National Laboratory Advanced Photon Source. Protein samples were freshly prepared using size exclusion chromatography (GE Life Sciences, Superdex 75 10/300 GL) in a buffer containing 50 mM Tris pH 7.5, 150 mM NaCl, 5 mM DTT, and 2 mM TCEP. Fractions were loaded immediately after elution without further manipulation. Buffer collected one column volume after protein elution from the column was used to record buffer data before and after each protein sample. SAXS data were acquired manually; protein samples were loaded, then gently refreshed with a syringe pump to prevent x-ray damage. A Pilatus 2M detector provided q-range coverage from 0.015 Å^-1^ to 1.0 Å^-1^. Wide-angle x-ray scattering data were acquired with a Pilatus 300k detector and had a q range of 0.93 – 2.9 Å^-1^. Calibration of the q-range calibration was performed with a silver behenate sample. Twenty sequential images were collected with 1 sec exposure time per image with each detector. Data were inspected for anomalous exposures and mean buffer data were subtracted from sample data using the WAXS water peak at q∼1.9 Å^-1^ as a subtraction control. Details about the SAXS data analysis can be found in the Supporting Information.

### 5.4 ENSEMBLE

ENSEMBLE 2.1^28^ was used to determine a subset of conformations from an initial pool of conformers created by the statistical coil generator TraDES. ^51,52^ All modules were given equal rank, and all other ENSEMBLE parameters were left at their default values.

To achieve a balance between the concerns of over-fitting (under-restraining) and under-fitting (over-restraining) we performed multiple independent ENSEMBLE calculations with 100 conformers, *N*_*conf*_ = 100, as suggested by Ref, ^54^ and averaged the results from independent ensemble calculation or combined them to form ensembles with larger numbers of conformers (e.g., *N*_*conf*_ = 500). To address the possibility that changing the ensemble size could affect the structural properties of the ensemble, or its agreement with experimental observables, we re-performed the Sic1 SAXS+PRE ensemble calculations, but varied the ensemble size, *N*_*conf*_ (details in the Supporting Information). The determination of polymer properties and the agreement with experimental observables is robust in a range of *N*_*conf*_ from ca. 50-100. Below *N*_*conf*_ ≈ 50, agreement with restraining data (SAXS and PRE) is worsened, and the ensembles do not agree with validating data (smFRET and CSs). Above *N*_*conf*_ ≈ 150, ensembles are in agreement with the experimental observables, though increased ensemble-to-ensemble variation suggests that replicates is insufficient to ensure convergence. Larger ensembles are calculated quicker (> 72 hours for *N*_*conf*_ = 20 vs ca. 1 hour for *N*_*conf*_ = 100). Ensembles with 100 conformers were chosen to minimize the computational cost per ensemble calculation, and ensemble-to-ensemble variation.

NMR data was obtained from BMRB accession numbers 16657 (Sic1) and 16659 (pSic1). ^26^ A total of 413 PRE restraints were used with a typical conservative upper- and lower-bound on PRE distance restraints of ±5 Å. ^53,74^ This tolerance was used in computing the *χ*^2^ metric for the PRE data. CSs were back-calculated using the SHIFTX calculator ^47^ and a total of 90 C_*α*_ CSs and 85 C_*β*_ CSs were used. The CS *χ*^2^ metric was computed using the experimental uncertainty *σ*_*exp*_ and the uncertainty in the SHIFTX calculator (*σ*_*SHIF T X*_ = 0.98 ppm for C_*α*_ CSs and *σ*_*SHIF T X*_ = 1.10 ppm for C_*β*_ CSs ^47^). CRYSOL ^30^ with default solvation parameters was used to predict the solution scattering from individual structures from their atomic coordinates. A total of 235 data points from *q* = 0.02 to *q* = 0.254 Å^-1^ were used in SAXS-restrained ensembles. The SAXS *χ*^2^ metric was computed using the experimental uncertainty in each data point.

Accessible volume (AV) simulations ^31,32^ were used to predict the sterically accessible space of the dye attached to each conformation via its flexible linker (Figure 1D). These calculations were performed using the AvTraj ^31^ v0.0.9 and MDTraj ^75^ v1.9.3 packages in Python 3.7.6. In the quasi-static approximation, the inter-dye distance dynamics within the AVs for a particular conformation are quasi-static on the timescale of the donor excited state (*τ*_*DA*_ *≤ τ*_*D*0_ = 3.7 ns). The per-conformer mean FRET efficiency is therefore *e* = ∫*E*(*r*_*DA*_)*P* (*r*_*DA*_)*dr*_*DA*_, where *P* (*r*_*DA*_) is the distribution of inter-dye distances resulting from the AV simulation for a particular conformation, and *E*(*r*_*DA*_) = (1 + (*r*_*DA*_*/R*_0_)^6^)^−1^. End-to-end distance reconfiguration times for IDPs and unfolded proteins are typically in the range 50-150 ns, ^76^ and so the end-to-end distance is also quasi-static on the timescale of *τ*_*DA*_. The back-calculated ensemble-averaged 〈*E*〉_*ens*_ is calculated as the linear average of the per-conformer FRET efficiencies 〈*E*〉_*ens*_ = *e*. The quasi-static approximation gives the same 〈*E*〉_*ens*_ within error as a more computationally demanding method which considers Monte-Carlo simulations of the photon emission process and Brownian motion simulations of dye translation diffusion within the accessible volume (detail in the Supporting Information). Further support for the quasistatic averaging approach used, comes from multiparameter *E* vs *τ*_*DA*_ histograms (Figure S2) which provide complementary information of inter-dye distances and dynamics, but with different experimental integration times.

The uncertainty in 〈*E*〉_*ens*_, *σ*_*E,ens*_, is ca. 0.01, which is a combination of SEM and uncertainty in *R*_0_. Differences 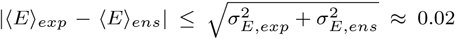 indicate no disagreement between back-calculated and experimental mean transfer efficiencies. A comprehensive description of the ENSEMBLE calculations, restraints and back-calculations can be found in the Supporting Information.

### 5.5 Polymer scaling analysis

The distance 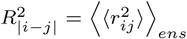 between C_*α*_ atoms is an average first over all pairs of residues that are separated by |*i* − *j*| residues, and then over all conformations in the ensemble. The apparent scaling exponent *v* was estimated by fitting an ISP calculated for each *N*_*conf*_ = 100 ensemble to the following expression:

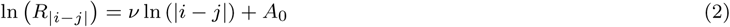

Eq. 2 is derived for homopolymers in the infinitely long chain limit. Following Peran and coworkers, ^37^ for finite-length chains, a lower bound of |*i* − *j*| > 15 was used to exclude deviations from infinitely-long-chain scaling behaviour at short sequence-separations and an upper bound of |*i* − *j* < |*n*_*res*_ − 5 was used to exclude deviations due to “dangling ends.” With these restrictions, finite-length homopolymers are expected to be well fit by Eq. 2. Evenly spaced points in log-log space were used during fitting. Fitting the entire 15 < |*i* − *j*| < *n*_*res*_ − 5 range was used to obtain *v. A*_0_ was either fixed at log(5.51) (*l*_*p*_=4 Å) or left as a free fitting parameter. To test for differences in scaling behaviour at intermediate and long sequence separations, the 15 < |*i* − *j* < |*n*_*res*_ − 5 range was evenly divided into intermediate *v*_*int*_ (15 *≤* |*i* − *j*| *≤* 51) and long *v*_*long*_ regimes (51 < |*i* − *j*| *≤ n*_*res*_ − 5).

## 6 Associated content

### 6.1 Supporting information

Extended description of smFRET and SAXS experiment/analysis, ENSEMBLE methods, and additional tables

## 7 Author information

### 7.2 Notes

The authors declare no competing financial interests

## 8 Acknowledgements

This work was supported by the Natural Sciences and Engineering Research Council of Canada (Grant No. RGPIN 2017–06030 to C.C.G. and Grant No. RGPIN-2016-06718 Fund 490974 to J.F.K). J.F-K., and T.H.G thank the National Institutes of Health for support under Grant 5R01GM127627-03. T.M. was supported by funding from St. Jude Children’s Research and the American Lebanese Syrian Associated Charities. We thank Dr. Taehyung Chris Lee for help preparing the Sic1 Y14A sample and S. Chakravarthy, J. Hopkins and all BioCAT beamline staff at the Advanced Photon Source for assistance with SAXS measurements. Use of the Advanced Photon Source was supported by the U.S. Department of Energy under contract DE-AC02-06CH11357.

