## Supplementary Information for "Integrating multiple experimental data to determine conformational ensembles of an intrinsically disordered protein"

#### Contents

**1 Single-molecule fluorescence spectroscopy** **S2**

1.1 Protein expression and purification . . . . . S2

1.2 Phosphorylation . . . . . S3

1.3 Labelling of proteins for FRET . . . . . S3

1.4 Single-molecule fluorescence instrumentation . . . . . S3

1.5 Single-molecule fluorescence sample conditions . . . . . S4

1.6 Burst search, filtering, calibration factors . . . . . S4

1.7 Förster radius . . . . . S5

1.8 Precision and accuracy of smFRET measurements . . . . . S5

1.9 Two-dimensional lifetime vs transfer efficiency plots . . . . . S6

1.10 Inferring distances from transfer efficiencies using polymer models . . . . . S6

**2 Small Angle X-ray Scattering experiments** **S7**

2.1 Model-free analysis (Guinier analysis, distance distribution function) . . . . . S7

2.2 SAXS analysis with explicitly represented flexible chains (ENSEMBLE,EOM, MFF) . . . . . S8

**3 Integrative structural modelling** **S8**

3.1 ENSEMBLE . . . . . S8

3.2 Ensemble size . . . . . . . . . . S9
3.3 PRE calculations . . . . . . . . . . S9
3.4 Chemical shifts calculations . . . . . . . . . . S10
3.5 SAXS calculations . . . . . . . . . . S11
3.6 Accessible volume FRET calculations . . . . . . . . . . S12
3.6.1 Averaging over conformations and accessible volumes . . . . . . . . . . S12
3.6.2 Comparison with physicochemically different dye pair . . . . . . . . . . S13

**4 Calculation of global polymeric dimensions** **S13**

**5 Supporting Tables** **S14**

**6 Supporting Figures** **S18**

### 1 Single-molecule fluorescence spectroscopy

#### 1.1 Protein expression and purification

A double cysteine variant of Sic1 (-1C-T90C) was generated via site directed mutagenesis. Sic1 -1C-T90C (henceforth Sic1) was expressed recombinantly as a Glutathione S-transferase (GST) fusion protein in *Escherichia coli* BL21 (DE3) codon plus cells grown in LB medium (LB Broth Lennox, Sigma L3022). 2 L cultures were grown in LB medium with 100 $\mu$g/ml carbenicillin until the OD600 reached ≈0.8-0.9. The cells were then induced with 1 mM IPTG overnight at 16 °C. The cells were harvested by centrifugation (3214×g, 60 min, 4 °C). After removing the supernatant pellets were either used for lysis immediately, or were frozen at -20 °C for future use in the near future.

The cell pellets were re-suspended in lysis buffer (PBS, 1 mM EDTA, 2 mM DTT, protease inhibitor cocktail (Roche)) and sonicated on ice for ≈ 5 minutes (30' on, 30' off). The resulting solution was then centrifuged (42.1 Rotor, Beckman Coulter, 30000 rpm, 60 min, 4 °C) to remove the insoluble fraction. The GST-Sic1 protein in the soluble fraction was purified by using glutathione-Sepharose affinity chromatography. The GST-Sic1 construct was incubated with the GST column (glutathione agarose resin, 786-280, G-Biosciences) for 2 hours at 4 °C. After thorough washing of the column to remove non-specifically bound proteins, the column buffer was changed to cleavage buffer (50 mM TRIS, pH 8.0, 0.5 mM EDTA, 1 mM DTT) and Sic1 was cleaved from the column with TEV protease (expressed recombinantly in *Escherichia coli*) overnight at 4 °C.

After digestion with TEV protease, the flow-through and wash from the GST-column predominantly contains Sic1 and TEV protease. The flow-through and wash are subjected to a final polishing step using cation-exchange chromatography (Bio-Scale Mini Macroprep High S 5 mL cartridge 732-4132, Biorad BiologicLP low pressure chromatography system). A typical ion exchange gradient optimized for Sic1 is 10% to 50% buffer B over 20 column volumes (100 mL) at a 1 mL/min flow rate. Buffer A is 50 mM Tris pH 7.5, 1mM EDTA, 2 mM DTT 50 mM NaCl, 0.001% Tween 20, while buffer B is identical but with higher ionic strength (1M NaCl). Tween 20 is present to prevent non-specific adsorption of the protein to the column and tubing of the chromatography system. A peak eluting at ≈ 40 mS/cm is identified as pure Sic1 by SDS-PAGE. Fractions containing only Sic1 (≈ 10 kDa) were pooled, concentrated, and buffer exchanged to either storage buffer (PBS buffer pH 7.4 with 3 M GdmCl, 2 mM DTT) and stored at -80°C or labelling buffer (see below). The correct molecular mass of the purified protein was verified by ESI-MS. The Y14A mutant Sic1 (-1C-T90C-Y14A) was generated via site directed mutagenesis from the aforementioned double-cysteine mutant and was expressed, purified and characterized using the same protocol.

##### 1.2 Phosphorylation

Phosphorylated samples were prepared by treatment of Sic1 with Cyclin A/Cdk2 (prepared according to Huang et al., [32])at a kinase:Sic1 ratio of 1:100 in the presence of 50 fold excess of ATP and 2.5 mM $MgCl_2$ overnight at 30 °C. The yield of phosphorylation reaction was determined by ESI-MS. Under these conditions the dominant species are 6- and 7-fold phosphorylated Sic1 (10195 Da and 10274 Da respectively) with a small fraction of 5-fold phosphorylated Sic1. After phosphorylation, the samples were buffer exchanged into PBS buffer pH 7.4 with 3 M GdmCl to prevent aggregation, denature kinase, and denature any phosphatases which may have inadvertently entered the solution. The samples were kept on ice in 4°C and measured within 24 hours.

##### 1.3 Labelling of proteins for FRET

The Sic1 smFRET construct with cysteines at positions -1C and 90C was labelled with Alexa Fluor 488 $C_5$ Maleimide (ThermoFisher Scientific, Invitrogen, A10254) and Alexa Fluor 647 $C_2$ Maleimide (ThermoFisher Scientific, Invitrogen, A20347). The protein was buffer exchanged and concentrated (Amicon Ultra-0.5 mL, UFC500396) to $\sim$ 200 $\mu M$ in degassed 50 mM PBS pH 7.0 3 M GdmCl and reduced using 10 mM DTT. The sample was incubated at room temperature to allow the cysteines to reduce. DTT was then removed by buffer exchange with 50 mM PBS pH 7.0 3 M GdmCl at 4°C and the protein concentration $\sim$ 200 $\mu M$ checked by the absorption at 280 nm. Alexa Fluor 647 $C_2$ Maleimide to the solution at a dye:protein ratio of 1.5:1 and the sample degassed by gently blowing with nitrogen for 1 minute. The solution was gently shaken at room temperature for 2-3 hours and the reaction stopped with with 5$\mu$L of BME ($\beta-$mercaptoethanol. The sample was then diluted into Buffer A (50 mM Tris pH 7.5, 1mM EDTA, 2 mM DTT 50 mM NaCl, 0.001% Tween 20) to further quench the reaction and prepare for ion exchange. Cation exchange (Bio-Scale Mini Macroprep High S 5 mL cartridge 732-4132, Biorad BiologicLP low pressure chromatography system) was used to separate the species with a single acceptor label from doubly labelled and unlabelled sample. Unreacted Alexa dyes (negatively charged) did not bind to the High S column. The single-labelled species sample was then concentrated and labelled with Alexa Fluor 488 $C_5$ Maleimide using the same procedure as for the acceptor. Cation exchange was again used to separate the doubly heterolabelled (donor and acceptor) from the acceptor only sample. The final FRET labelled sample was concentrated and buffer exchanged into PBS buffer pH 7.4 with 3 M GdmCl, 2 mM DTT and stored at -80 °C. The correct mass of the doubly labelled sample was confirmed by mass spectrometry.

##### 1.4 Single-molecule fluorescence instrumentation

Single-molecule fluorescence experiments were performed on a custom-built multiparameter confocal microscope equipped with a high numerical aperture (NA) infinity-corrected microscope objective (1.4 NA/100X Plan-Apochromat objective, 420790-9900, Carl Zeiss, Canada). The donor dye was excited with linearly polarized light centered at 480 nm using frequency doubled output from a Tsunami Ti:Sapphire (Spectra-Physics) laser in mode-locked operation with a pulse repetition rate of 80 MHz. The acceptor dye was excited with linearly polarized light from a continuous wave 635 nm diode laser (World StarTech, TECRL-25GC-635-TTL-A). To perform microsecond alternating laser excitation (ALEX)[36], the donor and acceptor excitation sources were alternated with "on" and "off" times $T_{donor} = T_{acceptor} = 50$ $\mu$s using an acousto-optic modulator (AOM) (Isomet, 1205C-2) for the donor excitation and direct TTL modulation for the acceptor excitation. Sample fluorescence was collected by the microscope objective, focused onto a 75 $\mu$m pinhole, and then separated into four channels first by separating according to wavelength range (donor or acceptor emission) using a dichroic filter (640DCXR, Chroma), and then by polarization using cube polarizing beam splitters. To additionally filter for Alexa Fluor 488 fluorescence, a bandpass filter BP530/50 and long pass filter E500LP (both Chroma) were used. To select for Alexa Fluor 647 fluorescence, the bandpass filter HQ685/80 and long pass filter HQ655LP (both Chroma) were used. Fluorescence photons were focused onto single-photon avalanche diodes (SPADs); either PDM-50CT SPADs (MPD, Italy) for the donor photons or COUNT-100C (Laser Components, USA) for acceptor photons. Photon arrival times relative to the start of the experiment (macrotimes) and relative to the exciting pulses (microtimes) were recorded using a PicoHarp 300 (PicoQuant, Germany) Time Correlated Single Photon Counting module and a router module (PHR800, PicoQuant, Germany). This instrumentation allows the simultaneous detection of the intensity, anisotropy, lifetime, and spectral properties of individual molecules.

#### 1.5 Single-molecule fluorescence sample conditions

Immediately prior to measurement samples were diluted to ~50 pM in either (*i*) PBS buffer: 10 mM sodium phosphate and 140 mM NaCl pH 7.0, 1mM EDTA (matches NMR measurements of Ref [46]) or (*ii*) Tris buffer: 50 mM Tris and 150 mM NaCl, pH 7.5. (matches SAXS measurements which were performed in high concentrations of Tris and reducing agents to scavenge radicals and prevent radiation damage). No difference in $\langle E \rangle_{exp}$ was detected when comparing buffer conditions and results are shown for Tris buffer conditions. Dilution of the smFRET samples from stock concentration in 3M GdmCl to single-molecule concentration results in approximately 60 nM residual concentration of GdmCl. Additionally, the SAXS measurements include 5 mM DTT, and 2 mM TCEP to scavenge radicals and prevent radiation damage but which are detrimental to fluorophore performance; while the smFRET measurements use 143 mM 2-mercaptoethanol (BME, 1:100 v/v dilution) and 5 mM 2-mercaptoethylamine (MEA) for photoprotection and increased brightness. The smFRET samples also contain 0.001 % Tween 20 for surface passivation.

The smFRET experiments were performed by exciting the donor dye with an average laser power of 140 $\mu$W (measured at the back aperture of the objective). For ALEX of the donor and acceptor, the power used for exciting the acceptor dye was adjusted to match the acceptor emission intensity to that of the donor (approximately 50 $\mu$W). A total sample volume of 30 $\mu$L was applied to a plasma cleaned coverslip and the confocal volume was chosen at 10 $\mu$m above the coverslip into solution. Sample evaporation was minimized to a point where it was not detected by sealing the top surface of the coverslip with a rubber spacer and a glass slide. Measurements were performed for $\approx$ 2 hours at the ambient temperature of the lab (20 °C).

#### 1.6 Burst search, filtering, calibration factors

The acquired data were subjected to multiparameter fluorescence analysis[40, 61] and ALEX filtering[36]. The burst search was performed using an All Photon Burst Search (APBS)[16, 48] with $M = 10$, $T = 500$ $\mu$s and $L = 50$. Bursts are identified as consecutive photons with intensity >20 kcps (20 times greater than background), that is, successive photons with inter-photon lag-time <50 $\mu$s. Only bursts with a total number of photons $L = 50$ after donor-excitation were kept for further analysis. Furthermore, selected bursts were required to have approximately symmetrical mean photon arrival times after donor and acceptor excitation $\left|T_X^{\mathrm{Dexc}} - T_X^{\mathrm{Aexc}}\right| < 300\ \mu s$ to filter out bursts corrupted by photobleaching or photoblinking[40]. Finally, the stoichiometry was required to be 0.25< S< 0.7 to select only molecules with both a photophysically active donor and acceptor fluorophore.

The transfer efficiencies of single molecules are obtained from $E = n_a/(n_a + \gamma n_d)$, where $n_d$ and $n_a$ are the numbers of donor and acceptor photons in each burst corrected for background, donor leakage into the acceptor channel, and acceptor direct excitation[28]. The $\gamma$ factor accounts for differences in the quantum yields of the dyes, and detection efficiencies. The $\gamma$ factor was determined using a 10 bp and 17bp dsDNA sample and fitting the relationship between the species $\langle E_{\mathrm{PR}}\rangle$ and $\langle S_{\mathrm{PR}}\rangle^{-1}$ as described in Ref [36]. The Sic1 specific $\gamma$ factor was found as $\gamma_{Sic1} = \gamma_{DNA}\frac{\phi_A^{Sic1}}{\phi_D^{Sic1}}\frac{\phi_D^{DNA}}{\phi_A^{DNA}}$. The overall $\gamma = 0.76 \pm 0.04$ was determined for all samples. Repeated measurements over a month indicate that the $\gamma$ factor is very stable on this timescale ($\gamma = 0.71 \pm 0.04$ after 1 month). The quantum yields of the donor and acceptor were determined based on the relative lifetime method ($\tau_X/\tau_Y = \phi_X/\phi_Y$). We found little variance of $\tau_{D0}$ and $\tau_A$ across samples (Table S1) suggesting that the sample specific dependent $\gamma$-factor does not change between samples (i.e., with phosphorylation or Y14A mutation).

The donor leakage was determined from the donor-only species which were selected from the two-dimensional $E$ vs $S$ histogram as described in Ref [36]. The average donor leakage factor $Lk \approx 0.014$ (Table S1) indicates that even for very bright bursts with $F_D^{\mathrm{Dexc}} = 100$ the number of photons which leak into the acceptor channel is minimal $F_{\mathrm{lk}} = Lk \cdot F_D^{\mathrm{Dexc}} \approx 1$ and does not significantly affect the determination of $\langle E\rangle$. Similarly, from the ratio of the absorption cross-sections of the donor and acceptor at the donor excitation wavelength $Dir \approx 0.005$ and so $F_{\mathrm{dir}} < 1$ even for very bright bursts.

#### 1.7 Förster radius

The Förster radius $R_0$ was calculated assuming a relative dipole orientation factor $\kappa^2 = 2/3$ and the refractive index of water $n = 1.33$. The overlap integral $J$ was measured for each sample and found not to change upon phosphorylation or Y14A mutation. As discussed above, there was minimal variation in $\tau_{D0}$ (Table S1) suggesting minimal variation in $\phi_D$. The $R_0$ was therefore calculated to be $R_0 = 52.2 \pm 1.1$ Å for all samples, and variation between samples within this uncertainty.

Subpopulation specific steady state anisotropies for the donor in the presence of the acceptor (i.e., the steady state anisotropy in the presence of $k_{FRET} > 0$, $r_{DA}$ - not be confused with inter-dye distance $r_{DA}$) were found to vary from 0.108 to 0.163 (Table S1). As a general rule[4, 11, 27, 27, 33], $r_{DA} < 0.2$ is taken to indicate that sufficient rotational averaging has taken place such that one can assume that the dynamic orientational average of $\kappa^2 = 2/3$ is a good approximation, as usually observed for IDPs and unfolded proteins.

#### 1.8 Precision and accuracy of smFRET measurements

The histogram of the transfer efficiency $E$ is fit to a Gaussian function to extract the mean transfer efficiency $\langle E\rangle$. The precision of $\langle E\rangle$ is found to be $\sim 0.005$ based on sequentially performed measurements of the same sample, or from bootstrap analysis[15] (randomly sampling the obtained bursts with replacement and reperforming analysis on these bootstrapped samples).

The accuracy of $\langle E\rangle$ measurements (due to variation in the instrumental and sample dependent calibration factors) was estimated to be $\sim 0.02$ from repeated measurements separated by longer than 1 month. Repeated measurements are often within 0.01 transfer units, however since measurements are performed on the same microscope and sample dependent calibration factors are estimated using the same protocol, we estimate the "true" uncertainty to be $\sim 0.02$ to allow for the fact that errors in the instrumental and sample dependent calibration factors may be correlated. An accuracy of $\sim 0.02$ was estimated for similar FRET measurements in the Schuler lab [30, 64] based on repeated measurements on different microscopes. Finally, a multi-laboratory benchmarking study[28] of the reproducibility and accuracy of FRET measurements provides an estimate of the accuracy to be between 0.02 and 0.05.

#### 1.9 Two-dimensional lifetime vs transfer efficiency plots

Multiparameter fluorescence allows us to excluded possible interfering artifacts, such as insufficient rotational averaging of the dyes which would be detected by high steady-state anisotropy values. Additionally, it can be used to perform a consistency check using two-dimensional histograms of $E$ calculated from intensities and the donor fluorescence lifetime.

For a fixed inter-dye distance $r$, $\tau_{DA}(r)/\tau_{D0} = (1-E(r))$, where $\tau_{DA}$ is the donor lifetime in the presence of the acceptor, $\tau_{D0}$ is the donor only lifetime, and $E(r) = (1 + (r/R_0)^6)^{-1}$. This describes a straight line (the static FRET line) in a 2D histogram of $E$ vs $\tau_{DA}/\tau_{D0}$ (black straight line in Figure S2). For IDPs and denatured proteins studied to date, end-to-end distance reconfiguration times are typically in the range of 50-150 ns [57, 63, 64] while inter-photon times are in the range 1-10 $\mu$s. These chains rapidly sample a broad distribution of distances $P(r)$ (see below for models) within the burst and between subsequent photon detection events.

The mean transfer efficiency $\langle E\rangle$ is therefore an averaged value over the distribution of distances sampled,

$$\langle E\rangle = \int_0^\infty E(r)P(r)dr \tag{1}$$

The average lifetime is,

$$\tau_{DA} = \int_0^\infty tI(t)dt / \int_0^\infty I(t)dt \tag{2}$$

with time-dependent fluorescence emission intensity,

$$I(t) = I_0 \int_0^\infty P(r)\exp\left(-t/\tau_{DA}(r)\right)dr \tag{3}$$

The expected and observed dependencies of $\tau_{DA}/\tau_{D0}$ and $E$ for (p)Sic1 and (p)Sic1 Y14A are summarized in the 2D histograms in Figure S2. In all samples the observed 2D histogram show populations which lie on the expected curve. Unaccounted for quenching of the dyes, incorrect $R_0$ or calibration factors, or an inappropriate assumptions about distance dynamics would tend to shift these populations off of the expected curve.

#### 1.10 Inferring distances from transfer efficiencies using polymer models

An estimate of the root-mean-squared end-to-end distance $R_{ee}$ can be made from $\langle E\rangle$ by assuming a homopolymer model distribution of distances $P(r_{D,A})$ between the donor and acceptor fluorophores which are attached by flexible linkers at the protein termini [26, 31],

$$\langle E \rangle = \int_0^\infty E(r_{D,A}) P(r_{D,A} | R_{D,A} dr_{D,A}) \tag{4}$$

where $r_{D,A}$ is the inter-dye distance and where,

$$E(r_{D,A}) = \frac{1}{1 + \left(\frac{r_{D,A}}{R_0}\right)^6} \tag{5}$$

Since dyes are attached to the protein via flexible linkers, the inferred distance is not directly $R_{ee}$, but rather a slightly larger distance $R_{D,A} = \langle r_{D,A}^2 \rangle^{1/2}$. The inferred $R_{D,A}$ is often rescaled to $R_{ee}$ by modelling the dyes and linkers as an additional number of residues in the chain such that,

$$R_{ee} = \left( \frac{n}{n + n_{dyes}} \right)^\nu R_{D,A} \tag{6}$$

Here, $n$ is the number of residues in the chain and the scaling exponent is $\nu = 0.5$ or $\nu \sim 0.6$ for the Gaussian chain (GC) model and Self-Avoiding Walk (SAW) model respectively. Two proposed values for $n_{dyes}$ are $n_{dyes} = 5 \pm 3$ [21] and $n_{dyes} \approx 9$ [69]. Using Eqs. 4 and 6, $R_{ee}$ was estimated using three models for $P(r_{D,A})$: the GC model, a renormalization-group-derived SAW model [49, 56] and a recently proposed semi-empirical model in which the scaling exponent $\nu$ (rather than $R_{D,A}$) is varied (rather than fixed to $\nu = 0.6$) to achieve agreement with $\langle E \rangle$ (the SAW-$\nu$ method) [70]. The results are summarized in Table S2.

To infer $R_g$ from $R_{ee}$ requires an additional assumption about the polymeric nature of system under study, namely $R_g^2 = R_{ee}^2/G$. For the GC model $G = 6$, while for the SAW model, $G \approx 6.25$ [41]. For the SAW-$\nu$ model, $G(\nu)$ is a function of the fitted solvent quality $\nu$ [70].

### 2 Small Angle X-ray Scattering experiments

#### 2.1 Model-free analysis (Guinier analysis, distance distribution function)

The root-mean-squared radius of gyration $R_g$ was first computed using the Guinier approximation ($I(q) = I(0) \exp\left(-q^2 R_g^2/3\right)$) via the AUTORG program from the ATSAS 2.8.4 suite of software[51], imposing that the maximum scattering vector $q_{max}$ satisfies an inequality involving $q_{max}R_g$ (Table S4). As was observed in Ref. [7] the inferred $R_g$ from the Guinier approximation is dependent on the upper bound of $q_{max}R_g$ used. The accepted range for folded proteins $q_{max}R_g \leq 1.3$ is expected to be more limited for unfolded proteins, since higher-order terms in the series expansion from which the Guinier approximation is derived are non-negligible for more extended conformations [7, 12, 56]. As $q_{max}R_g$ decreases, the inferred $R_g$ approaches the $R_g$ inferred using the full SAXS profile, e.g., $p(r)$ method and ensemble-based methods (*see below*). However, the uncertainty in $R_g$ also increases since the linear fit uses less data points. The upper limit $q_{max}R_g \leq 0.9$ was chosen based on the report by Ref [7] that an accuracy better than 0.5 Å for the IDPs and unfolded proteins studied was only obtained in this range.

Alternatively, $R_g$ may be estimated from the distance distribution function $p(r)$ obtained using the indirect Fourier transform of the regularized scattering curve[37] (Table S4). This estimation was performed using DATGNOM from the ATSAS 2.8.4 suite of software[51].

#### 2.2 SAXS analysis with explicitly represented flexible chains (ENSEMBLE,EOM, MFF)

In the Ensemble Optimization Method (EOM)[6, 66], an initial pool of conformations of the disordered protein is represented by random configurations of the $C_\alpha$ trace based on sequence information and either a "native" (*n-pool*) or "random" (*r-pool*) database of dihedral ($\phi$,$\psi$) angles. For disordered proteins, "native" angles are proposed to be a better representation [66] though we compared both angle databases which resulted in similar average properties of final selected ensembles. A genetic algorithm is used to select sub-ensembles from the initial pool on the basis of the agreement between the linear average of the CRYSOL-calculated SAXS profiles of individual conformers and the experimental data. The minimum ensemble is then selected as a result of 10,000 generations of optimization. We used EOM 2.0 from the ATSAS 2.8.4 suite of software[51] with default behaviour (except where noted above). The results of the EOM analysis are displayed in Table S5.

For a detailed description of the ENSEMBLE workflow see section3.1, and Ref [39]. For the analysis in this section, the only restraint for ENSEMBLE was the SAXS data, and ensembles of $N_{conf} = 100$ were generated. The results of this analysis are found in Table S5. To calculate $G$ in Table S5, $R_g$ was calculated from the $C_\alpha$ coordinates of the conformations.

Riback et al.,[53, 54] have introduced a procedure for fitting SAXS data by pre-generating ensembles of conformations with different properties (specifically, the strength and patterning of inter-residue attractions) and extracting dimensionless "molecular form factors" (MFFs). The properties of interest are then inferred from those of the ensemble whose MFF best fits the data. In the original MFF method (Ref [53], MFF in Table S6) homopolymer ensembles were pre-generated and the inferred properties are $R_g$, $R_{ee}$ and $\nu$. An updated MFF[54] generated heteropolymer ensembles with protein-like hydrophobic and polar (H/P) patterning (attractive and self-avoiding respectively). This model can be used in either a two free parameter ($R_g$ and $\nu$, MFF-het2 of Table S6) or three free parameter ($R_g$, $\nu$, and $\Delta\nu_{ends}$ MFF-het3 of Table S6). The three-parameter model allows for deviations in power-law scaling exponent of $R_{ij}$ at long sequence separations.

### 3 Integrative structural modelling

#### 3.1 ENSEMBLE

The ENSEMBLE method[39], begins with an initial pool of all-atom conformers derived either from the statistical coil generator TraDES [18, 19] or from previous MD simulations[5, 39]. We use the default statistical coil generation which generates four "flavours" of conformation in equal proportion: ALPHA, BETA, COIL and GOR3. For each flavour the TraDES-2[18, 19] program seq2trj creates a probability distribution for dihedral angles based on the following information supplied: predicted structure (H=helix, E=extended/sheet, C=coil), % probability of Helix, % probability of sheet and % probability of coil. Thus ALPHA is "H 60 20 20", BETA is "E 20 60 20", COIL is "C 0 0 100", while GOR3 uses the three-state GOR algorithm[23] to predict secondary structure from the sequence. These probability distributions, along with a Leonard-Jones type potential for self-avoidance, are use to create all-atom sequence specific protein conformations. ENSEMBLE then employs a switching Monte-Carlo algorithm embedded within a simulated annealing protocol to select a subset of $N_{conf}$ conformations (an ensemble) for which the back-calculated data fit all the available experimental data.

#### 3.2 Ensemble size

To achieve a balance between the concerns of over-fitting (under-restraining) and under-fitting (over-restraining) we performed multiple independent ENSEMBLE calculations with 100 conformers, $N_{conf} = 100$ and averaged the results from independent ensemble calculations or combined these ensembles to form ensembles with larger numbers of conformers (e.g., $N_{conf} = 500$). Structural features resulting from over-fitting (fitting the "noise" in the experimental data) should be averaged out in independent ENSEMBLE calculations, while rare states which are conserved in independently calculated ensembles (and thus have evidence from the data) should accumulate in weight when the ensembles are combined[44].

To address the possibility that changing the ensemble size could affect the structural properties of the ensemble, or its agreement with experimental observables, we re-performed the Sic1 SAXS+PRE ensemble calculations, but varied the ensemble size, $N_{conf}$ (Figure S3). The determination of polymer properties and the agreement with experimental observables is robust in a range of $N_{conf}$ from ca. 50-100. Below $N_{conf} \approx 50$, agreement with restraining data (SAXS and PRE) is worsened, and the ensembles do not agree with validating data (smFRET and CSs). Above $N_{conf} \approx 150$, ensembles are, on average, in agreement with the experimental observables, though increased ensemble-to-ensemble variation suggests that 5 replicates is insufficient to ensure convergence. Larger ensembles are calculated quicker ($>$ 72 hours for $N_{conf} = 20$ vs ca. 1 hour for $N_{conf} = 100$). Ensembles with 100 conformers were chosen to minimize the computational cost per ensemble calculation, and ensemble-to-ensemble variation.

#### 3.3 PRE calculations

The Sic1 and pSic1 PRE data was obtained from the BMRB (Biological Magnetic Resonance Data Bank) with accession numbers 16657 and 16659 respectively[46]. The PRE effects are from six single-cysteine Sic1/pSic1 mutants using a nitroxide spin label (MTSL) coupled to cysteine residues in positions -1, 21, 38, 64, 83, and 90. A residue-by-residue comparison of the agreement between the PRE restraints and back-calculated values is shown in Fig. S5 for Sic1 SAXS and SAXS+PRE ensembles, and in Fig. S6 for pSic1 SAXS and SAXS+PRE ensembles.

We use a simplified description of the PRE experimental data as distance restraints, which is a compromise between accuracy and computational cost, i.e., the Gillespie-Shortle approximation[24, 25]. In this approximation: (1) The MTSL spin-label is not explicitly modelled, and dynamic averaging over the experimental integration time is not considered[9] and (2) ensembles are restrained by average distance restraints $d = \langle r^{-6} \rangle^{-1/6}$, rather than back-calculated intensities, (3) a uniform correlation time of the electron-spin interaction vector $\tau_c$ for all residues, when it could vary depending on the local flexibility of the spin label and protein backbone. However, the significant decrease in computational cost makes this approach common in integrative methods[2, 8, 43, 44], including integration of NMR, SAXS, and smFRET[3]. Explicit modelling of the MTSL label[59] and/or its dynamics[68] for IDPs and unfolded proteins suggests that the decrease in accuracy is minimal, but the increase in computational time is significant.

Acknowledging the approximate character of this approach, generous error margins (±5 Å) are placed on the restraining distances $d$, and the PRE energy function is a flat bottom potential[22, 25]. This means that the contribution of a given distance restraint to the PRE energy function is zero if $d_{ij}^{res} - L < d_{ij}^{ens} < d_{ij}^{res} + U$, where $d_{ij}^{res}$ is the restraining distance calculated from the experimental PRE data, $d_{ij}^{ens}$ is the corresponding distance calculated from the ensembles molecular coordinates, and $L = U = 5$ Å are the lower- and upper-bound error margins.

The PRE effects were converted to electron-proton distances using,

$$d_{ij}^{res} = \left( \frac{K}{\Gamma_{ij}} \left( 4\tau_c + \frac{3\tau_c}{1 + (\omega_H \tau_c)^2} \right) \right)^{1/6} \tag{7}$$

where $K = 1.23 \times 10^{-32}$ cm$^6$s$^{-2}$ is a constant for a nitroxide radical, $\omega_H/2\pi = 500$ MHz is the Larmor frequency of the proton, and $\Gamma$ is the PRE value. A uniform correlation time of the electron-spin interaction vector $\tau_c = 2$ ns was assumed for all residues. Although it could vary throughout the chain, doubling $\tau_c$ results in an *average* shift of $d^{res}$ by approximately 1.4 Å with a maximum shift of 2 Å. As this is within the error tolerance of the flat bottom PRE potential it will result in no change in energy. The distance restraints were filtered by not including distance restraints when the fitting error from $\Gamma$ propagated to $d^{res}$ was greater than 10 Å, or if $d^{res} > 40$ Å. In total, $N_{PRE} = 413$ restraints were used.

The back-calculated distances are $d_{ij}^{ens} = \langle r_{ij}^{-6} \rangle^{-1/6}$ where $r_{ij}$ is the distance between the $C_\beta$ atom of the spin-labelled residue $i$ amide protons of residue $j$. The Sic1 ENSEMBLE models do not include the cysteine mutants, and so the PRE effects with single cysteine residues at position -1 (GLY) are modelled as arising the GLY $C_\alpha$ atom. For EPR measurements on two folded proteins, explicit modelling of the two MTSL spin-labels results in a mean absolute error of 4.4 Å, vs 6.1 Å when $C_\beta$ atoms are used to approximate the distances[1]. Given that our PRE measurements use a single label, are restrained with $\pm 5$ Å tolerances, and are averages over a large number of heterogeneous conformations (allowing for the possibility of cancellation of error), we do not expect that explicit modelling of MTSL would significantly change our results (as was found in Refs [59, 68]).

A reduced $\chi^2$ inspired metric was used as a measure of the agreement of the ensembles with the PRE restraints,

$$\chi^2 = N_{PRE}^{-1} \sum_{i=1}^{N_{PRE}} \left[ \frac{d_{ens}(i) - d_{res}(i)}{\sigma_{res}(i)} \right]^2 \tag{8}$$

where $d_{res}(i)$ is the $i$-th PRE distance restraint and $d_{ens}$ is the $\langle r^{-6} \rangle^{-1/6}$ distance for the corresponding pair of residues averaged over all the conformations in the ensemble. $\sigma_{res}$ is the restraint tolerance (upper-bound = lower-bound = 5 Å). As the weighted residuals in the summation are not expected to be standard normally distributed, and the degrees of freedom is not equal to the number of data points, this $\chi^2$ metric is not expected to be distributed according to the $\chi^2$ distribution, or have reduced $\chi^2 \sim 1$ for a good fit. Rather it should be interpreted merely as a normalized measure of the residuals. Note, this $\chi^2$ measure is not the same as the PRE energy in ENSEMBLE, which uses a flat-bottom potential.

The large PRE restraint tolerances could, in principle, affect the relative weights of the PRE and SAXS restraints. However, ENSEMBLE gives each experimental data type equal weight. To demonstrate this, we re-performed the Sic1 SAXS+PRE ensemble calculations, but decreased the PRE tolerance to $L = U = 2.5$ Å (Figure S4). Polymer properties and agreement with other experimental observables are highly similar for both sets of ensembles.

#### 3.4 Chemical shifts calculations

Chemical shift (CS) data was also obtained from BMRB accession numbers 16657 and 16659[46]. Fig. S7 shows the agreement of the $N_{conf} = 500$ Sic1 TraDES random coil, SAXS+PRE and SAXS+PRE+CS ensembles with the $C_\alpha$ and $C_\beta$ CSs. The three ensembles have similar $C_\alpha$ CS residuals, with most improvement (either indirectly by implementing SAXS+PRE restraints, or directly by implementing CS restraints) for pre-proline residues. For $C_\beta$ CSs, almost all residues in the coil and SAXS+PRE ensembles have residuals within the prediction errror of the SHIFTX calculator ($\sigma_{SHIFTX} = 0.98$ ppm for $C_\alpha$ CSs and $\sigma_{SHIFTX} = 1.10$ ppm for $C_\beta$ CSs[47]), however the residuals have a systematic offset. Implementing the CS restraints significantly reduces this offset, such that the residuals are distributed closer to zero. The similarity of $C_\alpha$ CS residuals for ensembles with different global properties (coil vs SAXS+PRE/SAXS+PRE+CS) and the difference in $C_\beta$ CS residuals for ensembles with similar global properties (SAXS+PRE vs SAXS+PRE+CS) demonstrates the decoupling of local and global structural properties for polymeric objects.

For the chemical shifts,

$$\chi^2 = N_{CS}^{-1} \sum_{i=1}^{N_{CS}} \frac{(CS_{ens}(i) - CS_{exp}(i))^2}{\sigma_{exp}^2(i) + \sigma_{SHIFTX}^2} \tag{9}$$

where $CS_{ens}$ and $CS_{exp}$ are the ensemble averaged back-calculated CSs and experimental CSs respectively. $\sigma_{exp}$ is the experimental error and $\sigma_{SHIFTX}$ is the SHIFTX prediction error. $C_\alpha$ and $C_\beta$ CS agreement are calculated separately. As noted in Ref. [10], this $\chi^2$ measure should merely be considered a normalized measure of the residuals, as the estimated prediction error for derived for folded proteins may overestimate that of IDPs, and normalization by $N_{CS}$ assumes completely independent data.

##### 3.5 SAXS calculations

CRYSOL[65] was used to predict the solution scattering from individual structures ($i(q)$) using the default adjustable parameters. The back-calculated SAXS profile is the ensemble average $I_{ens}(q) = \langle i(q) \rangle$. The ENSEMBLE SAXS energy term is a harmonic energy function with residuals weighted by the experimental uncertainty.

A reduced $\chi^2$ metric was calculated as,

$$\chi^2 = N_q^{-1} \sum_{i=1}^{N_q} \left[ \frac{A \cdot I_{ens}(q_i) - I_{exp}(q_i)}{\sigma_{exp}(q_i)} \right]^2 \tag{10}$$

where $I_{ens}$ is the back-calculated SAXS profile, $I_{exp}$ is the experimental SAXS profile and $\sigma_{exp}$ is its standard deviation. The overall normalization constant $A$ is adjusted to match the overall intensities of the model and data. $N_q$ is the number of scattering angle data points in the SAXS curve (235 points from $q = 0.02$ to $q = 0.254$ Å$^{-1}$). In principle, the calculation of $\chi^2$ should include an additional uncertainty due to the uncertainty in the back-calculation of $I_{ens}$, in particular from implicit hydration modelling[29]. Due to correlations in the data[67], the degrees of freedom may not be equal to $N_q$. For these reasons, the SAXS $\chi^2$ should not be assumed to have the same statistical properties as the canonical $\chi^2$.

Fig. S8 compares experimental SAXS curves and those predicted for the TraDES random coil and SAXS+PRE $N_{conf} = 500$ ensembles. For both Sic1 and pSic1 there is significant improvement in the agreement with the SAXS data over the TraDES random coil. Some minimal patterning in the weighted residuals for the pSic1 SAXS+PRE ensemble persists; however, no attempt was made to optimize the default solvation parameters in CRYSOL, which may not be optimal for disordered proteins[29, 50].

#### 3.6 Accessible volume FRET calculations

Accessible volume (AV) simulations[35, 60] were used to predict the sterically accessible space of the dye attached to each conformation via its flexible linker. These calculations were performed using the AvTraj[35] v0.0.9 and MDTraj[45] v1.9.3 packages in Python 3.7.6. The AV simulations are a purely geometrical search algorithm and no dye–dye or dye–chain interactions are taken into account apart from excluded volume. Table S7 provides the dye geometrical parameters used.

The flexible linkers produce a distribution of inter-dye distances, $r_{DA}$, over which the FRET distance dependence must be averaged, even for a single conformation with fixed end-to-end distance $r_{ee}$. The appropriate averaging over both $r_{DA}$ and $r_{ee}$ depends on the timescale of their respective motions relative to the donor excited state lifetime $\tau_{DA}$ ($\tau_{DA} \leq \tau_{D0} = 3.7$ ns) [17, 58]. To calculate the average FRET efficiency for a single conformation in the ensemble, we assume quasi-static distributions of inter-dye distances (see below). To calculate the ensemble average FRET efficiency, $\langle E \rangle_{ens}$ we assume a quasi-static distribution of end-to-end distances (i.e., the linear average of the per-conformer mean FRET efficiencies. We justify these assumptions below and show that explicitly modelling diffusion and the photon emission process gives nearly identical results (within the uncertainty of back-calculation and experiment) as the substantially faster static limit approximation (i.e., static $r_{DA}$ in accessible volumes and static $r_{ee}$ in ensembles).

##### 3.6.1 Averaging over conformations and accessible volumes

In the limit that inter-dye dynamics are much faster than $\tau_{DA}$, the "transfer-rate-weighted" dynamic average limit results[17, 58],

$$\langle E \rangle = \frac{\int \left(\frac{R_0}{r_{DA}}\right)^6 P(r_{DA}) dr_{DA}}{1 + \int \left(\frac{R_0}{r_{DA}}\right)^6 P(r_{DA}) dr_{DA}} \tag{11}$$

However, if the distances are quasi-static on the timescale of $\tau_{DA}$, the inter-dye distances are fixed at the (randomized) positions at the time of donor excitation. Consequently, in the quasi-static limit,

$$\langle E \rangle = \int E(r_{DA}) P(r_{DA}) dr_{DA} \tag{12}$$

For Sic1, it is expected that end-to-end distances are quasi-static on the timescale of $\tau_{DA}$. For IDPs and unfolded proteins studied to date, end-to-end reconfiguration times are typically in the range 50-150 ns [64]. We estimated the timescale of end-to-end distance dynamics in Sic1 to be approximately 35 ns by fitting the autocorrelation function of the end-to-end distance fluctuations from a simulation of Sic1 with the recently developed AMBER99SBdisp forcefield[55]. Finally, experimental evidence that justifies this assumption comes from the consistency between the expected and observed dependencies of $\tau_{DA}/\tau_{D0}$ and $E$ in Fig. S2.

Next, we turned to the treatment of averaging the distribution of inter-dye distances $r_{DA}$ resulting from the flexible dye linkers for a fixed $r_{ee}$. We performed Monte-Carlo simulations of the photon emission process and Brownian motion simulations of dye translational diffusion, adapting the approach in Ref. [17]. We used our measured donor-only lifetime $\tau_{D0}$ and Förster radius $R_0$. There is little information about the translational diffusion of dyes tethered to proteins in the literature[17]. We therefore use the range of translational diffusion coefficients determined by Peulen et al., $D_{AL488} \approx 0.9 - 10$ $\mathring{A}^2 ns^{-1}$ and $D_{AL647} \approx 0.4 - 5$ $\mathring{A}^2 ns^{-1}$ for Alexa 488 and Alexa 647 respectively, attached to DNA. We used the number of bursts, and the number of photons in each burst, from the Sic1 smFRET experiment, but determined whether each photon was a donor or acceptor photon based on the simulation.

In the smFRET experiment the inter-photon times and time between bursts are much longer than the end-to-end distance relaxation time. We therefore initiated each simulated photon from a randomly chosen conformation from the $N_{conf} = 500$ SAXS+PRE ensemble. Each fluorophore is initiated at a random point chosen from within its accessible volume for that conformation. At each time step $\Delta t = 0.001$ ps, we first decide if de-excitation has occurred with probability $p_1(t) = \Delta t k_{ET}(t) + k_{D0}$ at time $t$, where $k_{ET}(t) = k_{D0}(R_0/r_{DA}(t))^6$ and $k_{D0} = \tau_{D0}^{-1}$. If de-excitation occurs it was through energy transfer with probability $p_2(t) = k_{ET}(t)/(k_{ET}(t) + k_{D0})$, or donor emission with probability $1 - p_2(t)$ and the "identity" of the photon (acceptor or donor, respectively) is recorded, along with the excitation to de-excitation delay time, $t_D$. If the donor remains in the excited state, the donor and acceptor dye positions are updated using an Itô diffusion process, given by $d\mathbf{X} = \sigma d\mathbf{W}$ where $\mathbf{W}$ is a random vector of independent standard Wiener processes. The variance, $\sigma^2$, of the translational diffusion process is given by $6D_{dye}\Delta t$. If the dye tries to leave the AV, the proposed move is rejected, and a new move is drawn.

For each burst we calculate its FRET efficiency $E = N_A/N_{ph}$, where $N_A$ is the number of acceptor photons and $N_{ph}$ is the total number of photons in the burst. We also calculate the donor lifetime $\tau_{DA} = \langle t_D \rangle$. Finally, we calculate the respective averages over all simulated bursts, $\langle E \rangle_{sim}$ and $\langle \tau_{DA} \rangle_{sim}$. Table S8 summarizes the results of these simulations. The simulated $\langle E \rangle$ and $\langle \tau_{DA} \rangle$ are in good agreement with the experimental values. Explicitly modelling diffusion and the photon emission process gives nearly identical results (within the uncertainty of back-calculation and experiment) as the substantially faster static limit approximation (i.e., static $r_{DA}$ in accessible volume and static $r_{ee}$ in ensemble).

##### 3.6.2 Comparison with physicochemically different dye pair

While the consistency of the SAXS, NMR and FRET data suggests that labels do not affect the long-range or global structure of the Sic1 ensemble, and additional check is to measure FRET efficiencies for a different pair of dyes with different physicochemical properties. In a previous publication[42] we measured $\langle E \rangle_{exp}$ for Sic1 -1C90C in the same buffer conditions as the NMR measurements, but using the dye pair tetramethylrhodamine-5-Maleimeide (TMR) and Atto647N maleimide. In contrast to Alexa 488 which carries a -2 charge, TMR is neutrally charged, more hydrophobic and has a molecular weight approximately 66% of Alexa 488. In contrast to Alexa 647, which carries a -3 charge, Atto647N is cationic (+1 charge), more hydrophobic, and approximately 69% the molecular weight of Alexa 647.

We re-calculated the expected $\langle E \rangle_{ens}$ for the $N_{conf} = 500$ SAXS+PRE ensemble using the same approach as for the Alexa 488 and Alexa 647 dye pair, but with AVs calculated for TMR and Atto647N (Table S7) and the measured Förster radius $R_0 = 60 \pm 2$ Å. The resulting $\langle E \rangle_{ens} = 0.51 \pm 0.02$ agrees reasonably with the measured $\langle E \rangle_{exp} = 0.47 \pm 0.02$ (Figure 7D of Ref. [42]).

### 4 Calculation of global polymeric dimensions

The root mean-squared $R_g$ is calculated as the mean-squared sum over all $C\alpha$-$C\alpha$ distances,

$$R_g = \sqrt{\frac{1}{2n^2} \sum_{ij}^{n} \langle r_{ij}^2 \rangle} \tag{13}$$

The hydrodynamic radius, $R_h$ is calculated from $C\alpha$ coordinates using the Kirkwood-Riseman approach neglecting the free draining term[38],

$$\langle R_h^{-1} \rangle = \frac{1}{n^2} \sum_{i \neq j} \langle r_{\mathrm{ij}}^{-1} \rangle \tag{14}$$

$G = R_{ee}^2/R_g^2$ and $\rho = R_g/R_h$ are calculated from $R_g$ and $R_h$ using the conformations $C\alpha$ coordinates, consistent with their polymer theory interpretation. $A$ is calculated from the eigenvalues of the radius of gyration tensor of each conformation and then averaged over the ensemble to calculated $\langle A \rangle$ for each ensemble (as described in Ref [13]).

### 5 Supporting Tables

**Table S1:** Parameters determined and used in correction factors and calculations for smFRET data

| | $Lk$ | $\tau_{D0}$ (ns) | $\tau_A$ (ns) | $r_{DA}$ [a] |
|---|---|---|---|---|
| Sic1 | 0.0202 | 3.555 | 1.367 | 0.131 |
| pSic1 | 0.0153 | 3.582 | 1.355 | 0.135 |
| Sic1 Y14A | 0.0102 | 3.593 | 1.397 | 0.163 |
| pSic1 Y14A | 0.0089 | 3.656 | 1.422 | 0.139 |
| MEAN+/- STD | $0.0136 \pm 0.005$ | $3.60 \pm 0.04$ | $1.39 \pm 0.03$ | $0.14 \pm 0.01$ |

[a] The steady-state anisotropy of the donor, in the presence of energy transfer.

**Table S2:** Inferred $R_{ee}$ using polymer models and smFRET $\langle E \rangle$[a]

| | GC | | SAW | | SAW-$\nu$ | |
|---|---|---|---|---|---|---|
| $n_{dyes}$ | 5 | 9 | 5 | 9 | 5 | 9 |
| Sic1 | 65.4 | 64.0 | 62.5 | 60.9 | 63.8 | 62.5 |
| pSic1 | 71.6 | 70.1 | 67.8 | 66.1 | 68.9 | 67.5 |

[a] All dimensions are in angstroms. In addition to the uncertainty inherent in the choice of polymer model, the accuracy of $\langle E \rangle$ is $\pm 0.02$ which roughly corresponds to an uncertainty in the inferred $R_{ee}$ of $\pm 2$ Å or 3%. The precision, for measurements on the same day ($\leq 0.005$) corresponds to an uncertainty in inferred $R_{ee}$ of $\pm 0.4$ Å. The experimental transfer efficiencies were $\langle E \rangle = 0.42 \pm 0.02$ and $\langle E \rangle = 0.36 \pm 0.02$ for Sic1 and pSic1, respectively.

**Table S3:** Inferred $R_g$ using $R_{ee}$ from Table S2 and model dependent $G$ [a]

| | GC | | SAW | | SAW-$\nu$ | |
|---|---|---|---|---|---|---|
| $n_{dyes}$ | 5 | 9 | 5 | 9 | 5 | 9 |
| Sic1 | 26.7 | 26.1 | 25.0 | 24.3 | 26.8 | 26.4 |
| pSic1 | 29.2 | 28.6 | 27.1 | 26.4 | 28.6 | 28.1 |

[a] All dimensions are in angstroms. The accuracy of $\langle E \rangle$ of $\pm 0.02$ corresponds to an uncertainty in the inferred $R_g$ of $\pm 0.8$ Å. The uncertainty due to the precision in $\langle E \rangle$ of $\leq 0.005$, for measurements on the same day ($\leq$ 0.005) is approximately $\pm 0.2$ Å. The uncertainty due to the uncertainty in $R_0$ corresponds to approximately $\pm 1.6$ Å.

**Table S4:** Radii of gyration $R_g$ (in Å) inferred from the SAXS data using different conditions/methods.

| | | Guinier | | $P(r)$ |
|---|---|---|---|---|
| $q_{max}R_g$ | 0.9 | 1.1 | 1.3 | - |
| Sic1 | $30 \pm 4.1$ | $28.3 \pm 2$ | $28.6 \pm 1.4$ | 30.29 |
| pSic1 | $31.8 \pm 1.8$ | $30.7 \pm 2.2$ | $29.9 \pm 1.2$ | 32.18 |

**Table S5:** Global properties of the Sic1 and pSic1 ensembles using the SAXS data and either the EOM or ENSEMBLE method [a]

| | Sic1 | | | pSic1 | | |
|---|---|---|---|---|---|---|
| | ENSEMBLE | r-pool | n-pool | ENSEMBLE | r-pool | n-pool |
| $R_{ee}$ (Å) | $76.06 \pm 3.6$ | 79.45 | 80.69 | $75.86 \pm 3.9$ | 85.61 | 86.73 |
| $R_g^{hyd}$ (Å) | $30.62 \pm 0.87$ | 30.19 | 30.13 | $31.99 \pm 0.33$ | 31.99 | 31.91 |
| $G_{selection}$ | $6.39 \pm 0.24$ | 7.4 | 7.68 | $5.97 \pm 0.46$ | 7.63 | 7.87 |
| $G_{pool}$ | $6.37 \pm 0.51$ | 6.16 | 6.12 | $6.35 \pm 0.30$ | 6.15 | 6.13 |

[a] All dimensions are in angstroms. n-pool: EOM native initial pool, r-pool: EOM random initial pool. Only SAXS data was used to restrain the EOM/ENSEMBLE results in this table. $G_{pool}$ and $G_{selection}$ are calculated for the initial pool and final selected ensemble of conformers respectively. $G$ was calculated using the $C\alpha$ coordinate calculated $R_g$. The ensemble results are presented as the MEAN±STD for five independently calculated $N_{conf} = 100$ ensembles.

**Table S6:** Global properties of the Sic1 and pSic1 ensembles using the SAXS data and MFF methods

| | Sic1 | | | pSic1 | | |
|---|---|---|---|---|---|---|
| | MFF | MFF-het2 | MFF-het3 | MFF | MFF-het2 | MFF-het3 |
| $R_{ee}$ (Å) | $76.97 \pm 0.38$ | $77.3 \pm 0.37$ | $71.60 \pm 0.97$ | $79.97 \pm 0.31$ | $80.05 \pm 0.38$ | $71.13 \pm 0.91$ |
| $R_g$ (Å) | $30.63 \pm 0.15$ | $30.74 \pm 0.15$ | $31.61 \pm 0.43$ | $32.1 \pm 0.13$ | $32.13 \pm 0.15$ | $33.14 \pm 0.43$ |
| $G$ [a] | $6.31 \pm 0.05$ | $6.32 \pm 0.05$ | $5.1 \pm 0.1$ | $6.21 \pm 0.03$ | $6.21 \pm 0.04$ | $4.61 \pm 0.08$ |
| $\nu$ | $0.580 \pm 0.008$ | $0.584 \pm 0.011$ | $0.559 \pm 0.012$ | $0.547 \pm 0.005$ | $0.547 \pm 0.006$ | $0.527 \pm 0.007$ |
| $\Delta\nu_{ends}$ | NA | NA | $-0.172 \pm 0.081$ | NA | NA | $-0.236 \pm 0.095$ |

[a] $G$ is calculated assuming that the $R_g$ provided by MFF method is based on the $C\alpha$ coordinates of each residue. Uncertainty in $G$ by propagation of error neglecting the possible correlation in errors in $R_{ee}$ and $R_g$.

**Table S7:** Geometric details regarding the Accessible Volume simulations for dyes[a]

| | Donor (this study) Alexa Fluor 488 C5 maleimide | Acceptor (this study) Alexa Fluor 647 C2 maleimide | Donor (Ref. [42]) TMR-5-Maleimide | Acceptor (Ref. [42]) Atto647N maleimide |
|---|---|---|---|---|
| $L$ | 20.5 | 21.0 | 11.0 | 21.0 |
| $w$ | 4.5 | 4.5 | 4.5 | 4.5 |
| $R_{dye,1}$ | 5.0 | 11.0 | 5.0 | 7.15 |
| $R_{dye,2}$ | 4.5 | 4.7 | 4.5 | 4.5 |
| $R_{dye,3}$ | 1.5 | 1.5 | 1.5 | 1.5 |

[a] All dimensions in Angstroms. Geometric dye parameters from Ref [35]. Dyes are modelled as solid ellipsoids defined by three radii attached to a flexible linker with width $w$ and maximum extension $L$.

**Table S8:** Summary of Sic1 $N_{conf} = 500$ SAXS+PRE ensemble simulation with simulated dye motion[a]

| | $\langle E \rangle$ | $\langle \tau_{DA} \rangle$ (ns) |
|---|---|---|
| Static ($D = 0$) | 0.400 | 2.91 |
| $D_{min}$[b] | 0.417 | 2.89 |
| $D_{max}$[c] | 0.432 | 2.78 |
| Experiment | 0.42 | 2.95 |

[a] $\langle E \rangle_{ens}$ is calculated for $N_{conf} = 500$ ensembles. The uncertainty in $\langle E \rangle_{ens}$ is ca. 0.01, which is a combination of SEM and uncertainty in $R_0$. The uncertainty in $\langle E \rangle_{exp}$ is 0.02. The precision of $\langle \tau_{DA} \rangle$ is 0.02-0.03 ns.

[b] Using the lower-bound estimate for the translational diffusion coefficients in Ref. [52].

[c] Using the upper-bound estimate for the translational diffusion coefficients in Ref. [52].

**Table S9:** Comparison of back-calculated and experimental $\langle E \rangle$ for ensembles with different restraint contributions [a]

| | $\langle E \rangle$ | |
|---|---|---|
| | Sic1 | pSic1 |
| Experiment | 0.42 | 0.36 |
| Computation | | |
| TraDES RC | 0.305 | 0.309 |
| SAXS | 0.251 | 0.266 |
| PRE | 0.604 | 0.500 |
| SAXS+PRE | 0.400 | 0.338 |
| SAXS+PRE+CS | 0.421 | - |

[a] $\langle E \rangle_{ens}$ is calculated for $N_{conf} = 500$ ensembles. The uncertainty in $\langle E \rangle_{ens}$, $\sigma_{E,ens}$, is ca. 0.01, which is a combination of SEM and uncertainty in $R_0$. The uncertainty in $\langle E \rangle_{exp}$ is 0.02.

**Table S10:** Y14A mutation and phosphorylation smFRET results [a]

| | Construct | $\langle E\rangle$ | $R_{ee}$ (Å) | $\nu$ |
|---|---|---|---|---|
| Sic1 | -1C T90C | 0.424 | 63.8 | 0.52 |
| | -1C T90C Y14A | 0.395 | 66.3 | 0.53 |
| pSic1 | -1C T90C | 0.359 | 68.9 | 0.54 |
| | -1C T90C Y14A | 0.332 | 71.8 | 0.55 |

[a] The measurements were performed on the same day and sample dependent correction factors do not vary between samples (precision in $\langle E\rangle_{exp} < 0.005$). Additionally, $R_0$ does not vary between samples. $R_{ee}$ inferred using the SAW-$\nu$ method with $n_{dyes} = 5$. The precision of $\nu$ from the precision in $\langle E\rangle$ is $\sim 0.002$. The precision of $R_{ee}$ is ~0.4 Å.

**Table S11:** Global dimensions of the TraDES RC ensemble, SAXS-only ensemble, and SAXS+PRE ensembles[a]

| | | $R_{ee}$ (Å) | $R_g$ (Å) | $R_g^{hyd}$ (Å) | $R_h$ (Å) |
|---|---|---|---|---|---|
| Sic1 | TraDES RC | $70.33 \pm 3.54$ | $27.88 \pm 0.48$ | $28.97 \pm 0.46$ | $21.0 \pm 0.2$ |
| | SAXS-only | $76.06 \pm 3.6$ | $30.09 \pm 0.87$ | $30.62 \pm 0.87$ | $22.07 \pm 0.26$ |
| | SAXS+PRE | $61.92 \pm 1.84$ | $28.37 \pm 0.16$ | $29.24 \pm 0.14$ | $21.49 \pm 0.04$ |
| | SAXS+PRE+CS | $61.54 \pm 1.98$ | $28.58 \pm 0.33$ | $29.53 \pm 0.31$ | $21.50 \pm 0.13$ |
| pSic1 | TraDES RC | $70.35 \pm 2.45$ | $27.93 \pm 0.67$ | $29.20 \pm 0.64$ | $21.0 \pm 0.3$ |
| | SAXS-only | $75.86 \pm 3.9$ | $31.06 \pm 0.69$ | $31.99 \pm 0.33$ | $23.23 \pm 0.17$ |
| | SAXS+PRE | $68.65 \pm 1.76$ | $29.93 \pm 0.22$ | $30.73 \pm 0.17$ | $22.77 \pm 0.07$ |

[a] Reported values are the mean ± standard deviation of 5 independently calculated ensembles with 100 conformations ($N_{conf} = 100$). $R_g$ can be calculated directly from the $C_\alpha$ coordinates of each conformation ($r_g$ and $R_g = \langle r_g^2\rangle^{1/2}$) or taking into account the implicit hydration shell around each conformation $R_{g,hyd}$.

**Table S12:** Nominally universal polymer properties of the calculated ensembles[a]

| | | $G$ | $\rho$ | $\langle A\rangle$ | $\Delta A$ | $\Delta R_{ee}$ |
|---|---|---|---|---|---|---|
| Polymer Theory | EV ($n \to \infty$) | $6.254 \pm 0.004$ [b] | $\sim 1.59$ [c] | $0.431 \pm 0.002$ [d] | $0.442 \pm 0.004$ [d] | 0.374 |
| | EV ($n = 90 - 100$) | 6.32 [e-g] | 1.27–1.39 [g,h] | 0.4377 [f] | 0.437 [f] | - |
| | $\theta$-state ($n \to \infty$) | 6 [i] | $\sim 1.5$ [i] | 0.396 [d] | - | 0.422 |
| Sic1 | TraDES RC | $6.37 \pm 0.51$ | $1.33 \pm 0.01$ | $0.438 \pm 0.02$ | $0.438 \pm 0.02$ | $0.352 \pm 0.02$ |
| | SAXS-only | $6.39 \pm 0.24$ | $1.36 \pm 0.02$ | $0.470 \pm 0.02$ | $0.398 \pm 0.04$ | $0.329 \pm 0.02$ |
| | SAXS+PRE | $4.78 \pm 0.27$ | $1.320 \pm 0.007$ | $0.346 \pm 0.005$ | $0.454 \pm 0.02$ | $0.414 \pm 0.01$ |
| | SAXS+PRE+CS | $4.64 \pm 0.24$ | $1.330 \pm 0.008$ | $0.342 \pm 0.02$ | $0.472 \pm 0.02$ | $0.442 \pm 0.03$ |
| pSic1 | TraDES RC | $6.35 \pm 0.30$ | $1.33 \pm 0.02$ | $0.438 \pm 0.02$ | $0.432 \pm 0.03$ | $0.366 \pm 0.02$ |
| | SAXS-only | $5.97 \pm 0.46$ | $1.34 \pm 0.02$ | $0.418 \pm 0.02$ | $0.427 \pm 0.03$ | $0.354 \pm 0.04$ |
| | SAXS+PRE | $5.26 \pm 0.25$ | $1.314 \pm 0.005$ | $0.369 \pm 0.01$ | $0.428 \pm 0.02$ | $0.398 \pm 0.01$ |

[a] Reported values are the mean ± standard deviation of 5 independently calculated ensembles with 100 conformations ($N_{conf} = 100$). $G = R_{ee}^2/R_g^2$ and $\rho = R_g/R_h$ are calculated using the $C\alpha$ radius of gyration, consistent with their polymer theory interpretation. $A$ is calculated from the eigenvalues of the radius of gyration tensor of each conformation and then averaged over the ensemble to calculated $\langle A\rangle$ for each ensemble (as described in Ref [13]).
[b] Ref [41] and references therein.
[c] Ref [14] and references therein.
[d] Ref [34] and references therein.
[e] Ref [41] Monte Carlo SAWs $n = 100$
[f] Ref [34] Monte Carlo SAWs $n = 90$
[g] Ref [62] coarse-grained $C_\alpha$ bead model $n = 100$
[h] Ref [14] RG result with leading order correction for finite $n = 90$
[i] Ref [20]

### 6 Supporting Figures

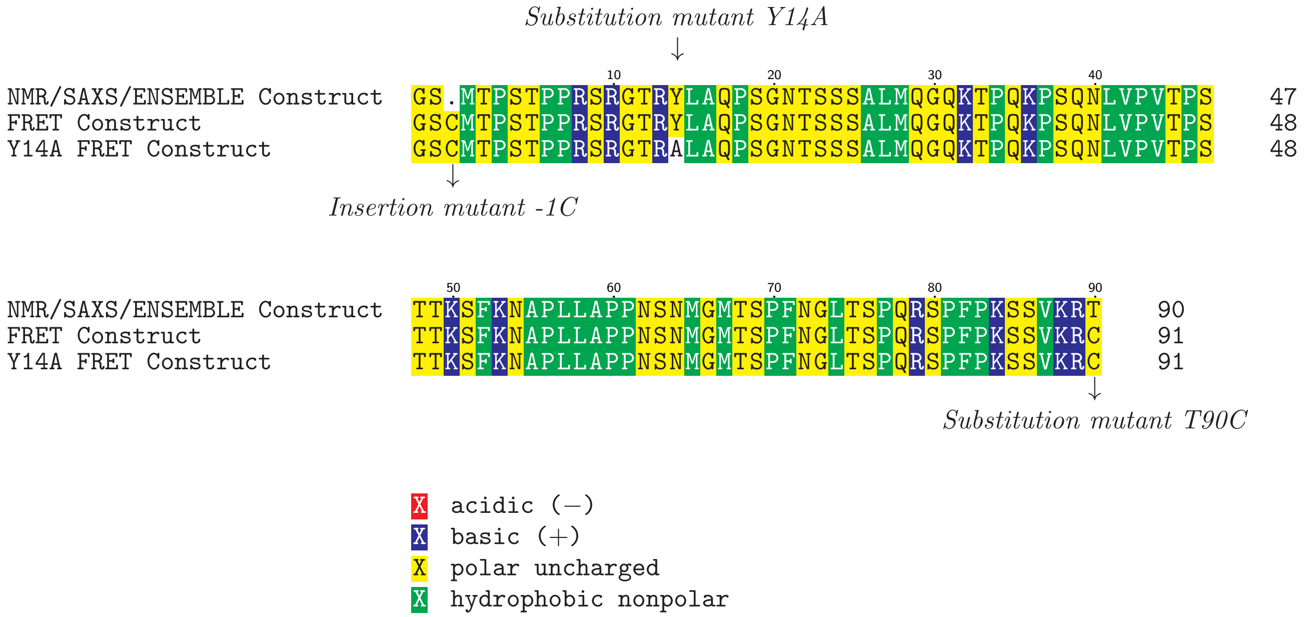


**Figure 1:** The main Sic1 N-terminal 1-90 constructs used in this paper .

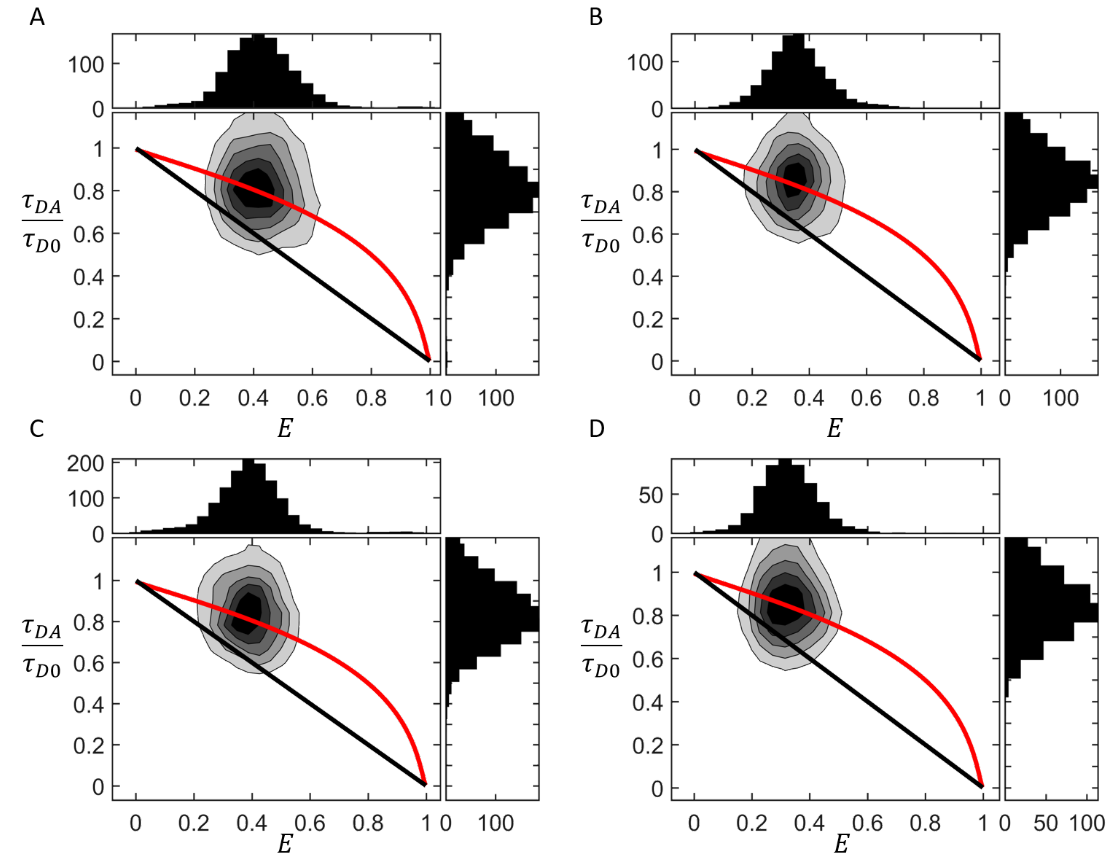


**Figure S2:** 2D-histograms of transfer efficiency versus relative fluorescence lifetimes $\tau_{DA}/\tau_{D0}$ for Sic1 (A), pSic1(B), Sic1 Y14A (C) and pSic1 Y14A (D). $\tau_{DA}$ and $\tau_{D0}$ are the donor lifetimes in the presence and absence of the acceptor, respectively. The lines within the plot represents the expected dependence of lifetimes on transfer efficiencies for a chain in which the inter-dye distance within the duration of a burst can be considered either fixed (black straight line) or averaged over the distribution of distances for a SAW $P(r_{D,A})$ (red curved line). Note: using a Gaussian chain or SAW-$\nu$ $P(r_{D,A})$ gives essentially overlaying curves.

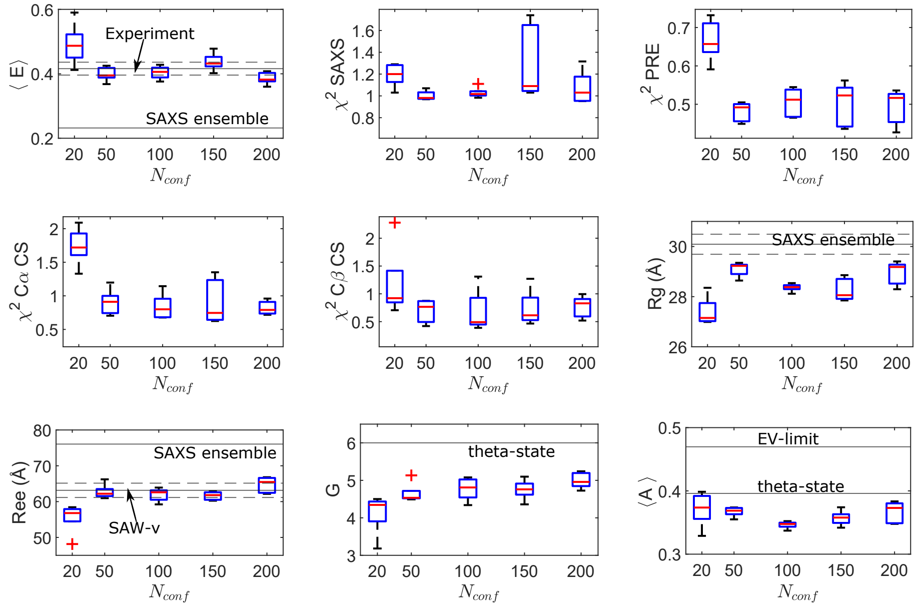


**Figure S3:** Polymer properties and agreement with experimental data used as restraints (SAXS+PRE) and validation (smFRET and CSs) for ensembles of varying size $N_{conf}$. Five replicates were performed for each $N_{conf}$ condition. Boxplot central mark indicates median, and edges indicate 25th and 75th percentiles. Whiskers extend to most extreme data points not considered outliers, with outliers marked by cross symbol. Solid horizontal lines show reference values as indicated, with dashed lines indicating an uncertainty range. Note: the agreement of the $N_{conf} = 500$ ensemble is better than the median of its five constituent $N_{conf} = 100$ as a result of cancellation of errors.

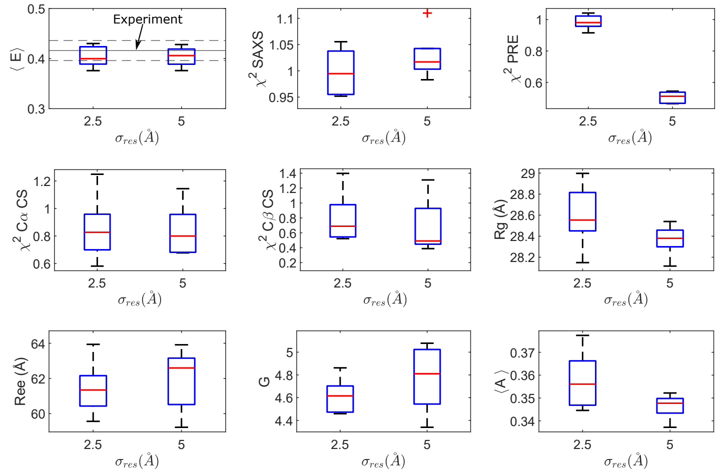


**Figure S4:** Polymer properties and agreement with experimental data used as restraints (SAXS+PRE) and validation (smFRET and CSs) for ensembles of varying PRE tolerance ($\sigma_{res}$). Five replicates were performed for each $\sigma_{res}$ condition. Boxplot central mark indicates median, and edges indicate 25th and 75th percentiles. Whiskers extend to most extreme data points not considered outliers, with outliers marked by cross symbol.

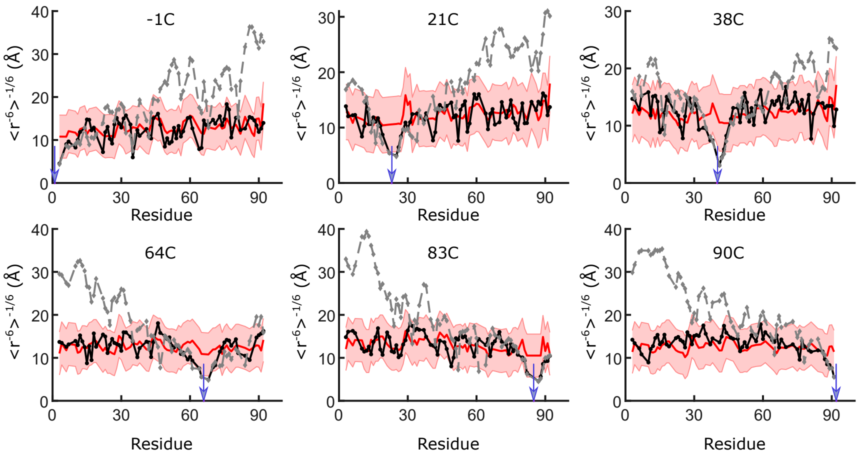


**Figure S5:** Sic1 experimentally derived PRE $\langle r^{-6}\rangle^{-1/6}$ distance restraints with ±5 Å tolerances (red shaded region) and back-calculated $\langle r^{-6}\rangle^{-1/6}$ distances for $N_{conf}$ = 500 Sic1 SAXS+PRE ensembles (black, solid lines) and Sic1 SAXS ensembles (grey, dashed lines). Distance restraints were derived from PRE effects measured for six different single-cysteine mutants spin-labelled at positions -1, 21, 38, 64, 83 and 90 (blue arrows)[46]. Back-calculated distances within the ±5 Å restraint tolerance (red shaded region) contribute no energy to the ENSEMBLE PRE energy function (flat-bottom potential).

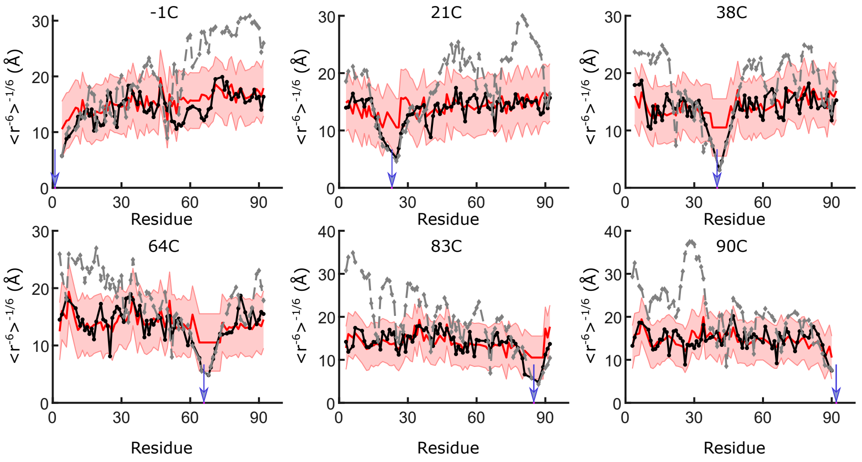


**Figure S6:** pSic1 experimentally derived PRE $\langle r^{-6}\rangle^{-1/6}$ distance restraints with ±5 Å tolerances (red shaded region) and back-calculated $\langle r^{-6}\rangle^{-1/6}$ distances for $N_{conf}$ = 500 pSic1 SAXS+PRE ensembles (black, solid lines) and pSic1 SAXS ensembles (grey, dashed lines). Distance restraints were derived from PRE effects measured for six different single-cysteine mutants spin-labelled at positions -1, 21, 38, 64, 83 and 90 (blue arrows)[46]. Back-calculated distances within the ±5 Å restraint tolerance (red shaded region) contribute no energy to the ENSEMBLE PRE energy function (flat-bottom potential).

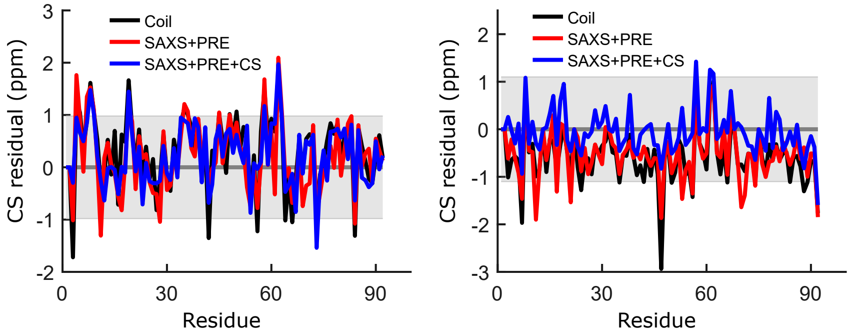


**Figure S7:** CS difference between the back-calculated and experimental CSs for the $N_{conf}$ = 500 Sic1 TraDES random coil, SAXS+PRE and SAXS+PRE+CS ensembles. The shaded grey region centred at zero indicates the reported prediction error of the SHIFTX calculator ($\sigma_{SHIFTX}$ = 0.98 ppm for $C_\alpha$ CSs and $\sigma_{SHIFTX}$ = 1.10 ppm for $C_\beta$ CSs[47]). Although for almost all residues, the difference lies within the prediction error, there is a systematic offset in $C_{beta}$ CSs in ensembles not restrained by CS data. However, and as expected, global and local properties of the ensembles are decoupled.

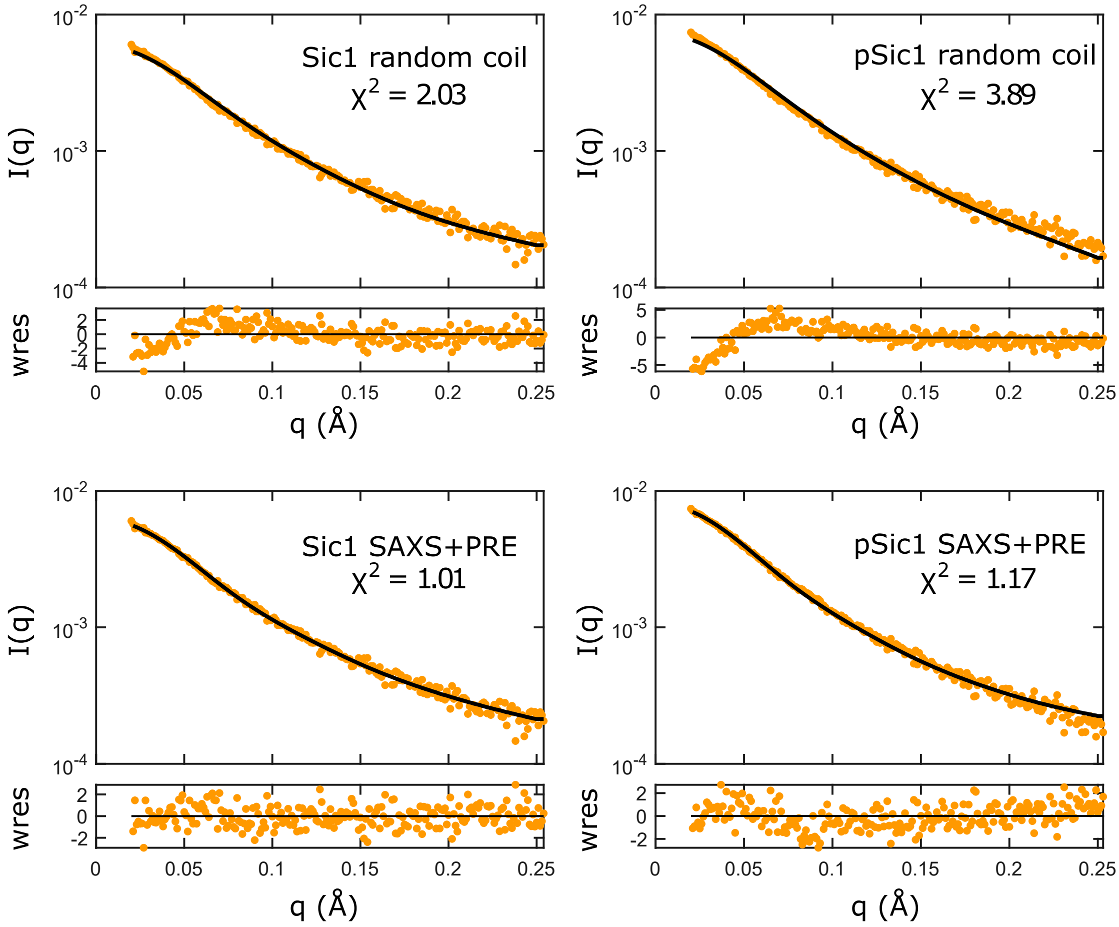


**Figure S8:** Comparison of back-calculated SAXS curves from example $N_{conf}$ = 500 ensembles with experimental data. The sub-panels show weighted residuals (wres).

[28] Björn Hellenkamp, Sonja Schmid, Olga Doroshenko, Oleg Opanasyuk, Ralf Kühnemuth, Soheila Rezaei Adariani, Benjamin Ambrose, Mikayel Aznauryan, Anders Barth, Victoria Birkedal, Mark E. Bowen, Hongtao Chen, Thorben Cordes, Tobias Eilert, Carel Fijen, Christian Gebhardt, Markus Götz, Giorgos Gouridis, Enrico Gratton, Taekjip Ha, Pengyu Hao, Christian A. Hanke, Andreas Hartmann, Jelle Hendrix, Lasse L. Hildebrandt, Verena Hirschfeld, Johannes Hohlbein, Boyang Hua, Christian G. Hübner, Eleni Kallis, Achillefs N. Kapanidis, Jae-Yeol Kim, Georg Krainer, Don C. Lamb, Nam Ki Lee, Edward A. Lemke, Brié Levesque, Marcia Levitus, James J. McCann, Nikolaus Naredi-Rainer, Daniel Nettels, Thuy Ngo, Ruoyi Qiu, Nicole C. Robb, Carlheinz Röcker, Hugo Sanabria, Michael Schlierf, Tim Schröder, Benjamin Schuler, Henning Seidel, Lisa Streit, Johann Thurn, Philip Tinnefeld, Swati Tyagi, Niels Vandenberk, Andrés Manuel Vera, Keith R. Weninger, Bettina Wünsch, Inna S. Yanez-Orozco, Jens Michaelis, Claus A. M. Seidel, Timothy D. Craggs, and Thorsten Hugel. Precision and Accuracy of Single-Molecule FRET Measurements—a Multi-Laboratory Benchmark Study. *Nature Methods*, 15 (9):669, September 2018. ISSN 1548-7105. doi: 10.1038/s41592-018-0085-0.

[52] Thomas-Otavio Peulen, Oleg Opanasyuk, and Claus A. M. Seidel. Combining Graphical and Analytical Methods with Molecular Simulations To Analyze Time-Resolved FRET Measurements of Labeled

Macromolecules Accurately. *The Journal of Physical Chemistry B*, 121(35):8211–8241, September 2017. ISSN 1520-6106. doi: 10.1021/acs.jpcb.7b03441.

[53] Joshua A. Riback, Micayla A. Bowman, Adam M. Zmyslowski, Catherine R. Knoverek, John M. Jumper, James R. Hinshaw, Emily B. Kaye, Karl F. Freed, Patricia L. Clark, and Tobin R. Sosnick. Innovative Scattering Analysis Shows That Hydrophobic Disordered Proteins Are Expanded in Water. *Science*, 358(6360):238–241, October 2017. ISSN 0036-8075, 1095-9203. doi: 10.1126/science.aan5774.

[54] Joshua A. Riback, Micayla A. Bowman, Adam M. Zmyslowski, Kevin W. Plaxco, Patricia L. Clark, and Tobin R. Sosnick. Commonly Used FRET Fluorophores Promote Collapse of an Otherwise Disordered Protein. *Proceedings of the National Academy of Sciences*, pages 8889–8894, April 2019. ISSN 0027-8424, 1091-6490. doi: 10.1073/pnas.1813038116.

[55] Paul Robustelli, Stefano Piana, and David E. Shaw. Developing a Molecular Dynamics Force Field for Both Folded and Disordered Protein States. *Proceedings of the National Academy of Sciences*, 115(21): E4758–E4766, May 2018. ISSN 0027-8424, 1091-6490. doi: 10.1073/pnas.1800690115.

[56] L Schäfer. *Excluded Volume Effects in Polymer Solutions as Explained by the Renormalization Group*. Springer, Berlin, 1999.
